# Human meningiomas arise from developmental meningeal mosaicism

**DOI:** 10.64898/2026.06.13.732068

**Authors:** Young-soo Chung, Hye Joung Cho, Taehyun Kim, Se Hoon Kim, Taek Chung, Seonah Choi, Tae-Hoon Roh, Ju Hyung Moon, Eui Hyun Kim, Chang Kyu Lee, Dong Ah Shin, Seong Yi, Yoon Ha, Keung Nyun Kim, Jong Hee Chang, Seok-Gu Kang, Sangwoo Kim

**Author notes:** These authors contributed equally to this work: Young-soo Chung, Hye Joung Cho. These authors jointly supervised this work: Seok-Gu Kang, Sangwoo Kim. Corresponding authors: Seok-Gu Kang: (82)-2228-2150, Sangwoo Kim: (82)-2228-2589.

## Abstract

Meningioma is the most common primary intracranial tumour, yet its genetic origin and the temporal sequence of mutational events remain poorly defined. Here, we analysed 80 triple-matched tumour, histologically normal meninges, and blood samples from 22 patients. *NF2* or *TRAF7* driver mutations are detectable in phenotypically normal meninges in 81.8% of cases (95% CI: 59.7–94.8%; VAF ∼0.02%), corroborated by high-depth sequencing, single-cell cloning, and phylogenetic analysis. By distinguishing developmental mosaic mutations from postnatal tumour-private mutations, we revealed distinct mutational signatures and resolved the temporal sequence of meningioma evolution. The developmental origin was further underscored in patients with multiple meningiomas, where identical driver mutations were shared across genomically distinct tumours and meninges, and in intraventricular meningiomas (IVM), where driver mutations were detected in distant cranial dura. Reconstruction of mutational timing revealed lineage-specific trajectories, linking mosaicism to diverse disease presentations including solitary meningioma, IVM, meningiomatosis, and *NF2*-related schwannomatosis. Together, these findings reveal an early origin of human meningioma, in which developmental mosaicism establishes a pre-neoplastic field within the meninges, providing a developmental framework for adult tumourigenesis.

## Main

Meningioma is a primary intracranial tumour arising from the proliferation of meningeal cells^1^. Although generally benign, its relatively high incidence (8–10 cases per 100,000 person-years)^2^, associated neurological symptoms, and the risk of growth or recurrence necessitate active clinical management, including wide surgical resection and rigorous recurrence risk stratification. Recent studies have explored the genomic and epigenomic profiles of meningioma, enabling clinically applicable molecular subgrouping^3–6^. While these classification systems successfully stratify tumours, they remain largely descriptive and fail to capture tumour dynamics. To fully define the mechanisms underlying tumour initiation and phenotypic diversity, elucidating the genetic underpinnings of driver mutations in anatomical and temporal context is essential^7–10^.

Large cohort genomic studies have identified two major molecular genotypes of meningioma, each comprising at least two genetic alterations: (i) *NF2*-type tumours, typically characterized by bi-allelic inactivation, typically involving translation-altering mutation combined with 22q loss; and (ii) non-*NF2* tumours, most commonly defined by a *TRAF7* mutation alongside a gain-of-function (GoF) mutation in either *AKT1* or *KLF4*^3^. However, the acquisition of multiple independent somatic events remains difficult to reconcile with low mutation burden (less than 1/Mb)^6^, relative genomic stability (e.g., few copy-number alterations)^3^, and the low proliferative activity expected for arachnoid cap cells, which are currently believed to be the cell of origin^11^. Moreover, classical models of stepwise somatic acquisition struggle to explain the development of multiple tumours, or meningiomatosis (MM), which may involve anatomically distant sites, including the spinal cord. While germline predisposition may contribute, in some cases, the presence of syndromic conditions such as *NF2*-related schwannomatosis (*NF2*-SWN, formerly known as neurofibromatosis type 2), caused by germline *NF2* mutations, complicates both the interpretation and diagnostic criteria^12^. These complexities imply a pathogenic framework of meningioma that incorporates anatomical sites, lesion burden, as well as timing of mutation acquisition.

### Patient cohort design and sample collection

To investigate the anatomical extent of driver mutations, we collected 80 triple-matched samples from 22 meningioma patients, comprising 32 tumours, 26 adjacent normal tissues (22 dura or falx, 2 cortex, and 2 lateral ventricular walls), and 22 peripheral blood samples (**Fig. 1a** and **Methods**). Patients were enrolled without selection for clinical, pathological, or molecular features to ensure cohort representativeness (**Methods**). Among them, 18 patients (P05–P18 and P21–P24) had solitary meningiomas (6 located in the convexity, 2 in the ventricular trigone, 8 in the skull base, and 2 in the spinal canal, **Extended Data Fig. 1a**). The remaining four patients (P01–04) presented with either concurrent or metachronous multiple meningiomas involving the brain and spine (**Extended Data Fig. 1b**). Specifically, five tumours were obtained from P01 and another five from P02, both of whom presented with multiple intracranial meningiomas. In P03, staged operation yielded one intracranial tumour and two spinal tumours. Although neither P02 nor P03 harboured germline *NF2* mutations or a family history, both met clinical criteria for *NF2*-SWN^13^ based on the presence of multiple meningiomas and vestibular schwannoma. In contrast, P01 and P04 exhibited multiple meningiomas without associated schwannomas and were therefore classified as meningiomatosis (MM).

**Figure 1.**
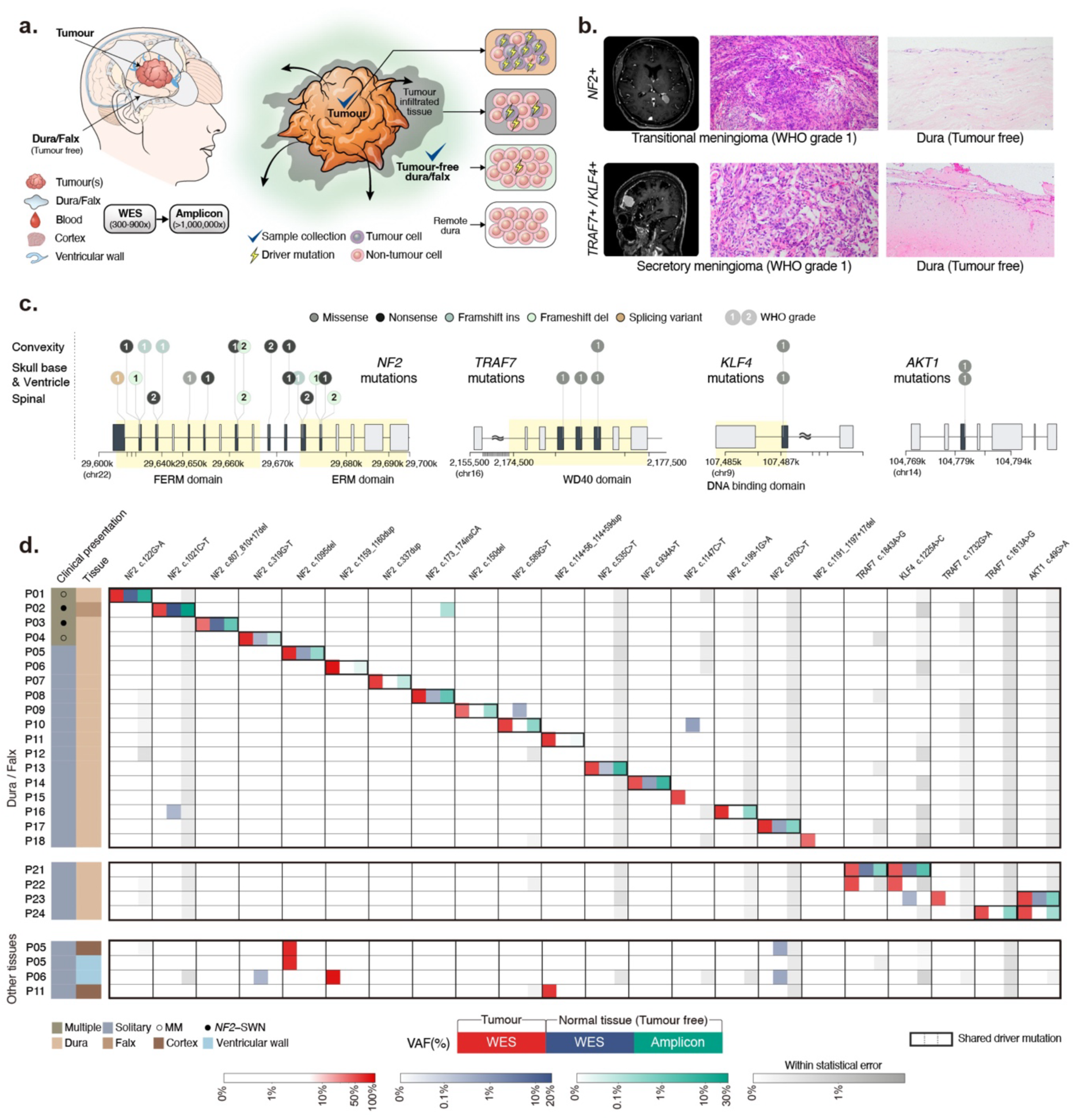
Identification of shared driver mutations in histologically normal meninges. **a,** Schematic of the surgical sampling strategy. Triple-matched tumour, histologically normal dura or falx, and peripheral blood were collected during craniotomy; cerebral cortex or ventricular wall was additionally obtained where clinically feasible. **b,** Representative MRI (left) and H&E staining of matched tumour (middle) and histologically normal dura (right) from an *NF2*-mutant (top) and a *TRAF7/KLF4*-mutant (bottom) case. High-power insets confirm the absence of neoplastic infiltration in dural tissue. **c,** Lollipop plots showing somatic driver mutations mapped to genomic coordinates and annotated protein domains. **d,** Driver mutation VAFs in matched samples per patient: tumour (WES; red), histologically normal meninges (WES; blue), and normal meninges by amplicon sequencing (AS; green). Grey gradients indicate sample-specific background noise thresholds derived from a binomial null distribution (see **Methods**). In 18 out of 22 patients (81.8%; 95% CI:59.7–94.8%), tumour-initiating driver variants were identified within matched normal meninges. **VAF**, variant allele frequency; **WES**, whole-exome sequencing; **AS**, amplicon sequencing.

Adjacent normal tissues were sampled at locations maximally distant from the tumour within surgical constraints to minimize tumour contamination. Normal dura was obtained from the edge of the craniotomy window, sufficiently distant from the tumour epicentre to avoid inclusion of the dural tail. In spinal meningiomas, dura was collected at the dorsal midline dural incision line to avoid iatrogenic spinal canal narrowing or cerebrospinal fluid leakage. The falx, cortex, and ventricular wall were sampled only when exposed within the surgical corridor during tumour access. Non-dural meningeal layers, including the arachnoid membrane, could not be obtained in sufficient quantities for sequencing analysis. All normal tissues were confirmed as histologically tumour-free by neuropathological examination, with no evidence of tumour infiltration (**Fig. 1b** and **Methods**).

### Presence of driver mutations in normal meninges

We performed whole-exome sequencing (WES; average coverage of 128x) on 32 tumour samples and matched peripheral blood to identify somatic mutations (**Methods**). *NF2* alterations were detected in 28 tumours (87.5%) across 18 patients (81.8%). All *NF2*-mutant tumours exhibited clear evidence of biallelic inactivation, comprising a deleterious mutation (nonsense, frameshift, splice-site variant, or structural variation) accompanied by chromosome 22q loss (**Fig. 1c**). Mutation sites were patient-specific (17 unique variants) and distributed throughout the coding region of *NF2*. In one patient (P12), who harboured 22q loss without an identifiable deleterious *NF2* mutation, long-read sequencing identified a cryptic structural variation (SV) resulting in the complete absence of wild-type transcripts. (**Extended Data Fig. 2**). Arm-level copy-number analysis further stratified tumours into molecular subgroups according to established criteria: MenG A (n = 3), MenG B (n = 27) and MenG C (n = 2) (**Extended Data Fig. 3a**). Tumours harbouring *NF2* mutation or 22q loss were distributed throughout the craniospinal axis, including the convexity, falx, skull base, ventricular trigone, and spinal canal. The remaining four patients (18.2%), whose tumours were located in the anterior region (that is, frontal convexity, frontal base, and sphenoid wing), harboured mutations in the WD40 domain of *TRAF7*; two additionally carried *KLF4* p.K409Q, and the other two had *AKT1* p.E17K mutations, both of which are known drivers in non-*NF2* meningiomas (**Fig. 1c**)^3^. No other recurrent meningioma-associated mutations were identified^14^. Overall, the mutational composition, low mutational burden (0.2–5.6/Mb), and anatomical specificity were consistent with previous reports^6^ (**Extended Data Fig. 4**).

We next examined adjacent normal tissues. High depth WES (average coverage of 336x), combined with variant calling strategies optimized for low-allele fractions (**Methods** and **Extended Data Fig. 5**), identified low-frequency somatic mutations (variant allele frequency (VAF), 0.2%–19.2%) in the dura or falx of 11 patients. These mutations occurred at identical genomic positions and involved the same base substitutions as those observed in the matched tumours (**Fig. 1d**). We further performed amplicon sequencing (AS; average coverage of 1,077,216x) targeting the 22 unique tumour mutation sites across all available tumour and meningeal tissues, except for the case harbouring SV (P12), for which targeted amplicon design was not feasible (**Methods**). All meningeal driver mutations were again detected in the AS with consistent VAF relative to WES (Pearson’s *R* = 0.67 ; p < 0.0001). In addition, AS identified eight further tumour-matched mutations in seven individuals at very low VAFs (0.02%–0.4%), all of which were exclusively present at the sites of matched tumour mutations and were significantly deviated from the background allele frequencies (**Methods** and **Extended Data Fig. 6a–b)**. Overall, 18 of 22 patients (81.8%), including 83.3% (15 out of 18) *NF2*-type and 75% (3 out of 4) non-*NF2* type cases, exhibited shared driver mutations in tumour-free meninges.

### Mosaic origin of tumour-normal shared driver mutations

To distinguish pre-existing developmental mosaicism from tumour infiltration as the source of the shared driver mutations, we analysed the profile of passenger (non-driver) mutations across matched tumour–meningeal pairs (**Fig. 2a**). If the mutations detected in adjacent meninges primarily reflect infiltration, dura or falx would mirror a substantial fraction of tumour-derived passenger mutations. By contrast, under a mosaic model, only a limited number of ancestral mutations would be shared, whereas most subsequently acquired tumour-associated mutations would remain private to one lineage.

**Figure 2.**
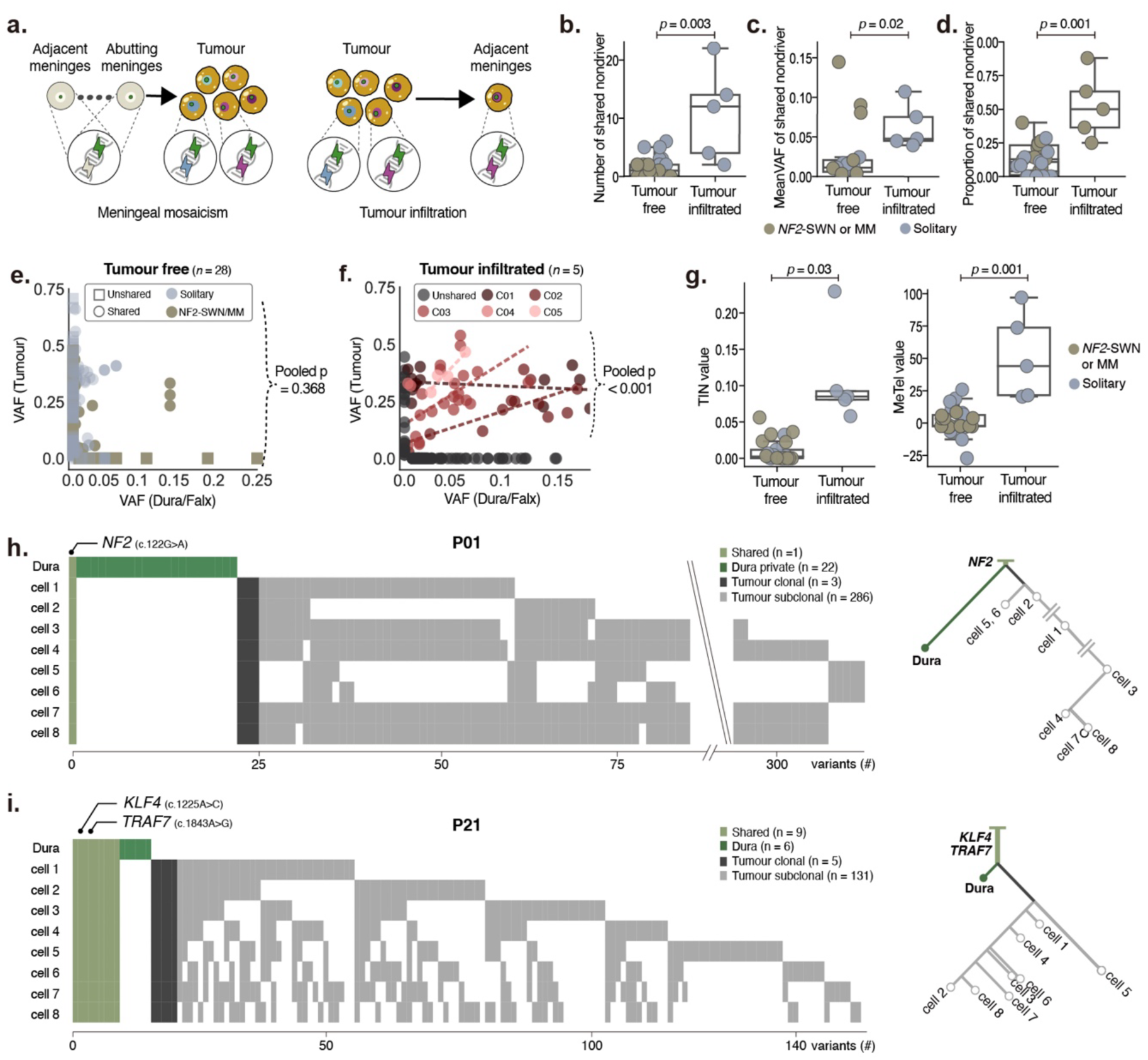
Shared driver mutations in tumour-free meninges originate from developmental mosaicism, not tumour infiltration. **a,** Schematic of two competing hypotheses for shared driver mutations in tumour and adjacent meninges: developmental mosaicism versus tumour infiltration. **b–d,** Comparison of shared mutation burden (**b**), VAF (**c**), and proportion of shared non-driver mutations (**d**) between tumour-free and pathologically confirmed infiltrated pairs (Mann–Whitney tests; *P* values shown). **e,** Non-driver mutation sharing between tumour and normal dura or falx across 28 tumour-free pairs. No significant positive VAF correlation was observed (fixed-effect meta-analysis of Fisher z-transformed correlations; pooled one-sided *P* = 0.368). **f,** Same analysis in five pathologically confirmed infiltrated controls (C01–C05), showing extensive non-driver mutation sharing and positive VAF correlation between tumour and matched dura or falx (*P* < 0.001). **g,** *In silico* tumour contamination estimates by TINC (left) and MeTel (right) for tumour-free and infiltrated meningeal pairs; values near zero indicate absence of clonal dissemination. **h, i,** Phylogenetic trees reconstructed from WES of single-cell-derived clones from P01 (**h**) and P21 (**i**). Mutation profiles (left) and phylogenetic trees (right) reconstructed from WES of single-cell-derived clones from P01 (h) and P21 (i). The x-axis represents the number of somatic variants per sample. Double diagonal lines in h indicate axis truncation. Tumour clones and matched meninges diverge from a common ancestor, with tumour-associated mutations predominantly acquired after lineage separation. **VAF**, variant allele frequency; **WES**, whole-exome sequencing.

Across 28 tumour–dura/falx pairs from 18 individuals harbouring shared driver mutations in tumour-free meninges, we identified 534 somatic mutations, including 31 drivers and 503 passengers (**Supplementary Table 1**). Among passenger mutations, only 42 (8.3%; mean, 1.5 per pair) were shared between tumour and dura/falx, whereas 307 (61.0%; mean, 11.0 per pair) and 154 (30.6%; mean, 5.5 per pair) were private to tumour and dura/falx, respectively. In 10 of the 28 pairs, no passenger mutations were shared.

To determine whether this pattern differed from *bona fide* infiltration, we analysed five additional tumour-dura/falx pairs with pathologically confirmed meningeal infiltration (C01–05; **Extended Data Fig. 1**) using the same sequencing and analytical workflow. Tumour-infiltrated pairs shared a substantially higher number of passenger mutations (mean 10.8 versus 1.5 per pair, Mann–Whitney *P* = 0.003) compared to the tumour-free pairs (**Fig. 2b**), and exhibited higher VAFs (mean VAF, 6.3% versus 2.7%, Mann–Whitney *P* = 0.019) (**Fig. 2c**). Notably, the three tumour-free pairs with relatively high shared-mutation VAFs all originated from patients with multiple meningiomas (*NF2*-SWN or MM), consistent with earlier mosaic acquisition. Likewise, the proportion of shared passenger mutations among all tumour somatic mutations was higher in infiltrated pairs (0.52 versus 0.10, *P* = 0.001) (**Fig. 2d**). Furthermore, in infiltrated cases, VAFs in tumour and matched dura/falx were positively correlated (fixed-effect meta-analysis of Fisher z-transformed correlations, pooled one-sided *P* < 0.01), indicating that the mutational profile in dura/falx largely mirrored that of the tumour (**Fig. 2e)**. By contrast, no such positive correlation was observed in tumour-free pairs (pooled *P* = 0.37) (**Fig. 2f)**. Finally, in silico estimates of tumour contamination using TINC^15^ clearly separated tumour-free from infiltrated dura/falx (**Fig. 2g, left**; 0.11 versus 0.01, Mann–Whitney *P* = 0.03), with values near zero indicating no detectable contamination, and MeTel^16^ analysis indicated a markedly higher likelihood of clonal dissemination in infiltrated samples than in tumour-free pairs (**Fig. 2g, right**; 51.1 versus 0.50; *P* = 0.001), with values near zero supporting independent origins. Together, these orthogonal analyses indicate that tumour infiltration contributes minimally to the shared driver mutations observed in tumour-free meninges.

We next examined clonal relationships directly at the single-cell level in two informative cases for which single-cell isolation and clonal expansion were available. In P01, an *NF2*-mutant case presenting with meningiomatosis, eight single-cell-derived tumour clones exhibited substantial genomic heterogeneity and shared only the *NF2* mutation with the corresponding dura (**Fig. 2h, left**). In P21, a solitary meningioma harbouring *TRAF7* and *KLF4* mutations, eight clones shared a limited set of mutations including *TRAF7* and *KLF4*, whereas most mutations were private either to the tumour clones or to the dura (**Fig. 2i, left**). Phylogenetic reconstruction in both cases placed the tumour clones and matched dura on diverging branches descending from a common ancestor, with most tumour-associated mutations acquired after lineage separation (**Fig. 2h, i, right**). These lineage relationships are most consistent with ancestral meningeal mutations followed by independent mutation accumulation, with any low-level tumour infiltration at the single-cell level insufficient to explain the observed mutational patterns.

Taken together, these observations support a developmental meningeal mosaic origin of the shared low-VAF driver mutations detected in tumour-free meninges and identify developmental mosaicism as a previously underappreciated mechanism contributing to meningioma initiation.

### Early developmental meningeal mosaicism gives rise to spatially distinct meningiomas

Having established a developmental mosaic origin, we next analysed the clonality and distribution of somatic mutations across matched tumour-dura/falx pairs to infer when these mutations arise during development. Mutations shared between tumours and dura/falx were predominantly clonal, with a high mean cancer cell fraction (CCF) of 86.0%, indicating acquisition before tumour–meningeal lineage divergence (**Fig. 3a**, left). By contrast, tumour-private mutations showed a bimodal distribution, comprising both clonal (mean CCF = 92.4%) and subclonal (mean CCF = 20.3%) subsets, consistent with acquisition after this early shared phase and during subsequent tumour evolution (**Fig. 3a**, right). Shared clonal mutations were enriched for C>T transitions (**Fig. 3b**) and SBS1 and SBS5 signatures, reflecting clock-like mutational processes associated with early postzygotic acquisition (**Fig. 3c**)^17^, whereas subclonal tumour-private mutations were enriched for SBS15 (15.7%), in line with mutational profiles reported in prior meningioma cohorts^3^. These data indicate that the earliest detectable mutations in meningioma arise during early development, preceding tumour formation and lineage divergence, and are reflected in the spatial distribution of tumours.

**Figure 3.**
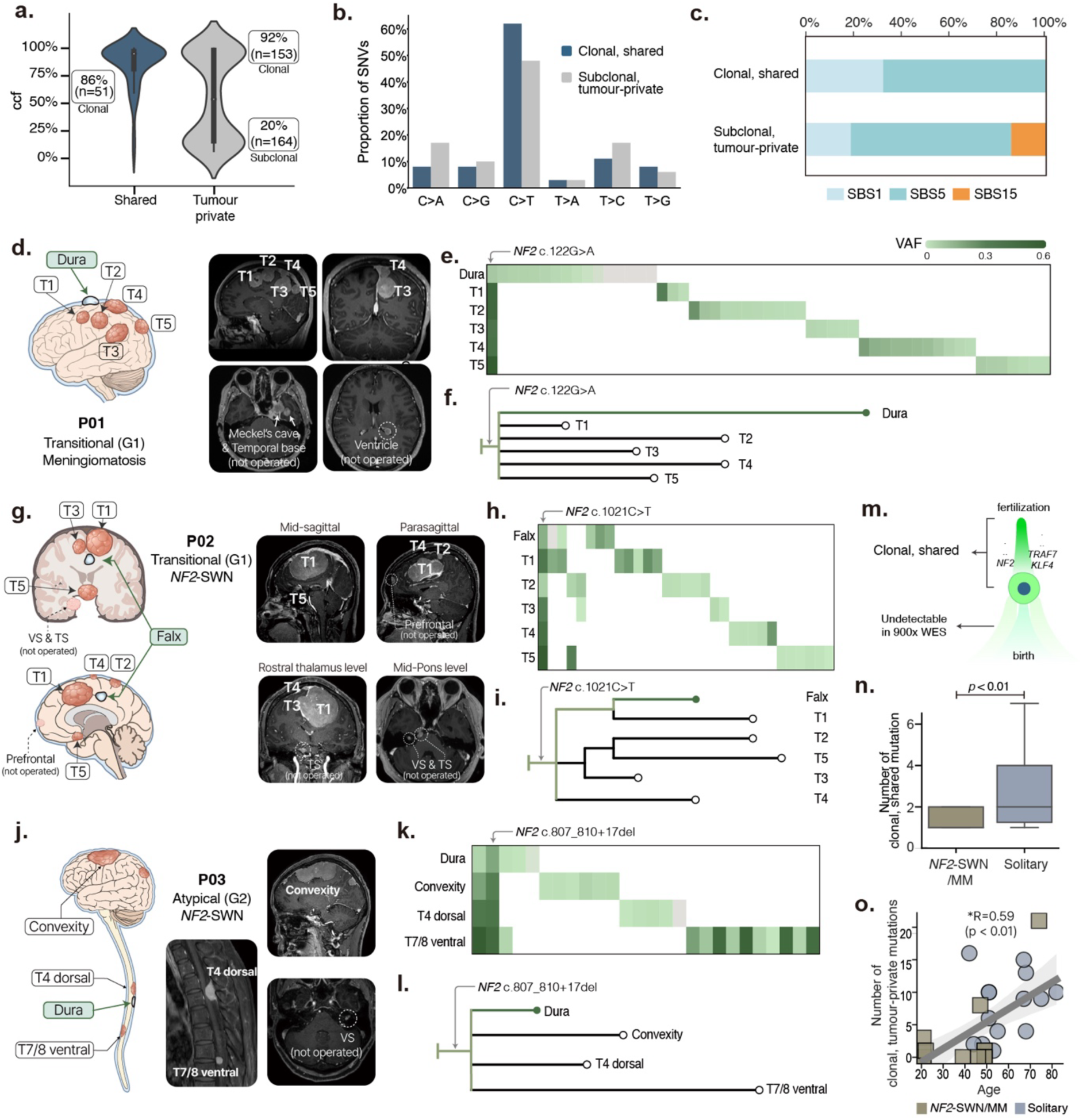
Developmental meningeal mosaicism underlies spatially distinct meningiomas. **a,** CCFs of shared (left; mean CCF = 86.0%) and tumour-private (right; clonal mean CCF = 92.4%, subclonal mean CCF = 20.3%) mutations. **b, c,** Base-substitution profiles **(b)** and COSMIC mutational signature compositions **(c)** of shared clonal versus tumour-private subclonal mutations. **d–l,** Multi-regional sampling, mutational profile, and phylogenetic reconstruction for P01 (MM; **d–f**), P02 (*NF2*-SWN; **g–i**), and P03 (*NF2*-SWN; **j–l**). In P01 and P02, five spatially separated tumours each share only the *NF2* driver mutation with adjacent dura or falx; in P03, synchronous spinal and metachronous intracranial tumours share only the *NF2* driver despite otherwise divergent mutational profiles. **m,** Schematic of the developmental mosaicism model, in which earlier acquisition corresponds to fewer clonal shared mutations. **n,** Shared clonal mutation burden in *NF2*-SWN or MM versus solitary meningioma pairs (1.4 vs. 2.9; t-test, *P* < 0.01). **o,** Correlation between tumour-private clonal mutation count and patient age (*R* = 0.59, *P* < 0.01). **CCF,** cancer cell fraction; **MM,** meningiomatosis; ***NF2*-SWN,** *NF2*-related schwannomatosis.

Early developmental meningeal mosaicism was evident in patients with multiple tumours, including both *NF2*-SWN and MM. In P01 (**Fig. 3d**), sequencing of five spatially distinct synchronous intracranial meningiomas revealed marked genomic heterogeneity, with only the *NF2* driver mutation (c.122G>A) shared across all tumours and the adjacent dura (**Fig. 3e**). Phylogenetic reconstruction placed the dura on the ancestral branch, supporting a model of independent tumourigenesis rather than clonal dissemination (**Fig. 3f**). This pattern was also observed in clinically diagnosed *NF2*-SWN cases. In P02, who presented with multiple tumours involving the falx, parasagittal convexity, and tuberculum sella (**Fig. 3g**), the *NF2* driver mutation (c.1021C>T) was consistently detected across all tumours and in the falx (**Fig. 3h–i**). Given the shared neural crest origin of midline meningeal structures and cranial nerve–associated tissues, these findings are consistent with mutation acquisition during early neural crest development. In P03, two synchronous spinal meningiomas (T4 dorsal and T7/8 ventral) and one metachronous intracranial meningioma (Convexity) exhibited genomic divergence, sharing only two ancestral mutations, including the *NF2* driver mutation (c.319G>T) (**Fig. 3j–l**). This further supports independent tumour formation from a shared mosaic origin in the meninges. In all three cases, the VAFs of dural or falcine *NF2* mutations were elevated (8.5% in P01, 18.8% in P02, and 1.3% in P03) relative to the cohort average of 0.58%. Notably, these individuals showed lesions across a broad anatomical spectrum, including an intraventricular meningioma (P01) and vestibular or trigeminal schwannomas (P02 and P03), supporting the widespread anatomical consequence of early *NF2* mosaicism.

We extended the mutational time frame to encompass the zygote-to-adult trajectory and aligned genetic events accordingly (**Extended Data Fig. 7**). Conceptually, clonal shared mutations may reflect mutational events acquired early in development, potentially during the prenatal period. Accordingly, when mosaicism arises very early, the number of clonal shared mutations would be expected to remain small, reflecting only a limited ancestral mutational burden before lineage divergence (**Fig. 3m**). Consistent with this hypothesis, tumour–dura/falx pairs from patients with *NF2*-SWN or MM harboured significantly fewer clonal shared mutations than those from patients with solitary meningiomas (1.4 vs. 2.9, t-test *P* < 0.01; **Fig. 3n**). This observation further supports a developmental framework for meningioma pathogenesis. We next focused on tumour-private clonal mutations. As clonal tumour mutations often reflect the genetic state of the cell of origin^18^, tumour-private clonal mutations were interpreted as lineage-restricted events acquired after divergence between tumour and meningeal lineages, likely during postnatal life (**Extended Data Fig. 7**). Indeed, the number of tumour-private clonal mutations correlated with patient age (*R* = 0.59, *P* < 0.01), consistent with age-associated somatic accumulation in normal cells (**Fig. 3o**). These observations define a model in which early postzygotic mosaic mutations establish a pre-neoplastic meningeal field, followed by the accumulation of lineage-restricted clonal mutations during tumour development.

### Developmental mosaicism and meningioma formation at anatomically distinct sites

Intraventricular meningiomas (IVMs) arise at anatomically distinct sites despite the absence of an arachnoid layer lining the ventricular walls, posing a challenge to current models of meningioma origin. The tela choroidea, a loosely organized connective tissue derived from the pia mater and situated at the ventricular interface, has been proposed as a potential origin site for IVMs, although this remains controversial.

From two patients with IVMs (P05 and P06), tumours located in the trigone of the lateral ventricle were resected, alongside samples from the LV roof (ventricular wall), parieto-occipital dura, and normal cortex acquired during initial corticectomy (**Fig. 4a–b**; see **Methods**); due to the risk of damaging the choroid plexus, tela choroidea tissue could not be obtained. WES of tumours revealed *NF2* frameshift indels (a 1-bp deletion in P05 and a 2-bp insertion in P06).

**Figure 4.**
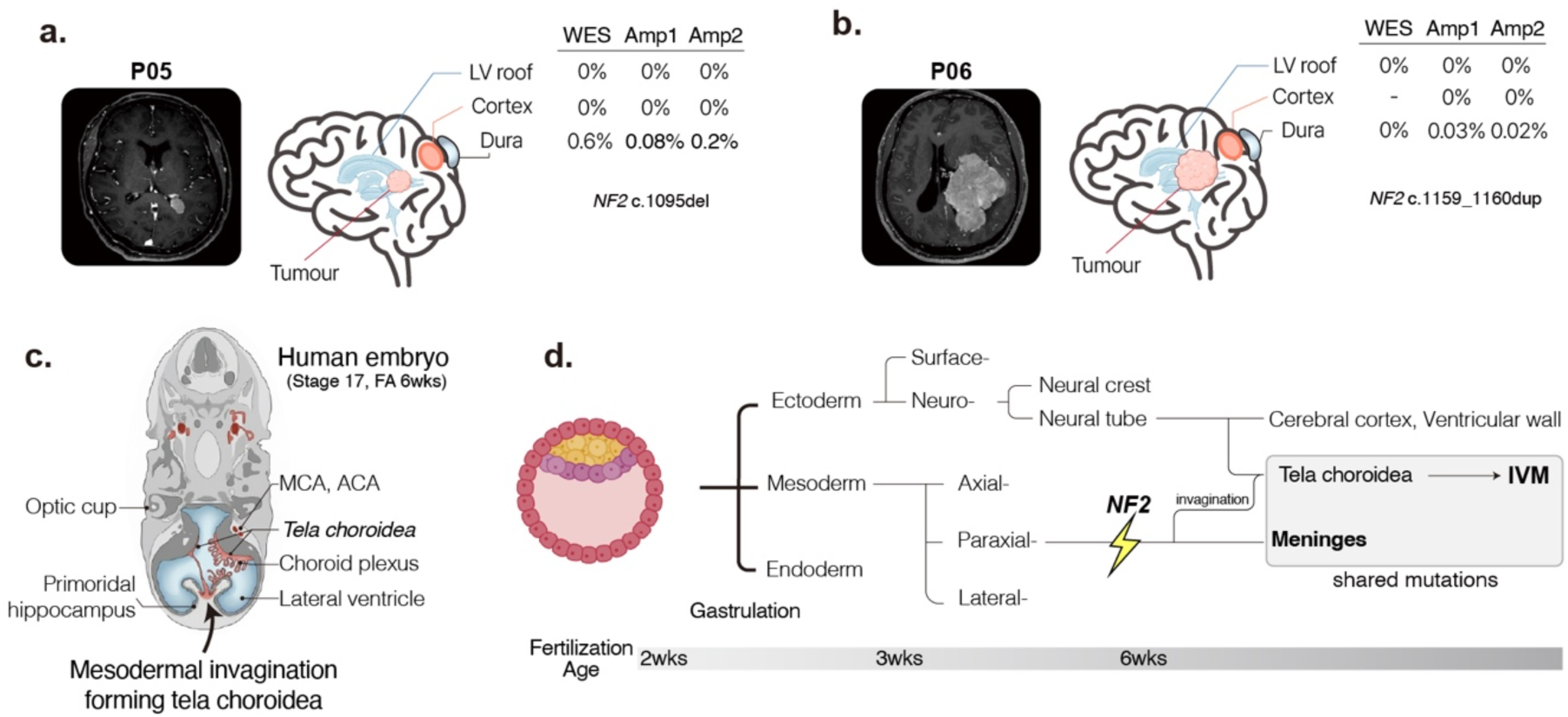
Intraventricular meningioma as a manifestation of early meningeal mosaicism. **a, b,** Representative IVM cases from P05 **(a)** and P06 **(b)**, showing MRI, sampling schematic, and driver mutation VAFs across matched tissues. *NF2* frameshift indels are detected in parieto-occipital dura (VAF 0.02–0.3%) but not in the lateral ventricular wall or cortex. **c,** Schematic of human embryological development at approximately 6 weeks of gestation (Carnegie stage 17), illustrating mesenchymal invagination and formation of the tela choroidea and choroid plexus. **d,** Reconstructed developmental lineage placing *NF2* mutation acquisition prior to embryological separation of ventricular and cranial meningeal compartments. **IVM**, intraventricular meningioma; **FA**, fertilization age; **VAF**, variant allele frequency.

High-depth AS of non-tumour tissues revealed the matching *NF2* frameshift indels at low VAFs (0.02%–0.3%) in the dura, but not in the LV roof or normal cortex. This discordance between the mutation-sharing pattern and anatomical proximity, given that the parieto-occipital dura is spatially distant and separated by intervening cortex and ventricular wall, challenges models invoking either a local origin from the ventricular wall or centrifugal tumour cell migration to the dura. Instead, the shared developmental lineage between the tela choroidea and meninges, in contrast to the distinct neuroectodermal origin of the ventricular root and cortex^19,20^, supports developmental *NF2* mosaicism within a mesenchymal compartment as a parsimonious explanation. These findings support early acquisition of *NF2* mosaicism during meningeal or related mesenchymal development, potentially before the anatomical divergence of ventricular and cranial meningeal structures, which is thought to occur around 6–7 weeks after fertilization (**Fig. 4c,d**). Together, these observations indicate that lineage-specific developmental mosaicism contributes to the anatomical localization of meningioma.

### Early developmental acquisition of major mutational events

The developmental mosaicism framework raises the possibility that key mutational events in meningioma may occur substantially earlier than clinical tumour presentation. Previous models have proposed that meningioma development proceeds through anywhere from the classical two-hit mechanism to more complex multi-step processes involving additional genetic alterations^21^, yet the temporal ordering of these events remains poorly defined. To investigate the timing of chromosomal aberrations, we performed single-nucleus DNA sequencing (Tapestri platform; see **Methods**) on dura or falx samples from three *NF2*-related meningiomas (**Fig. 5a**).

**Figure 5.**
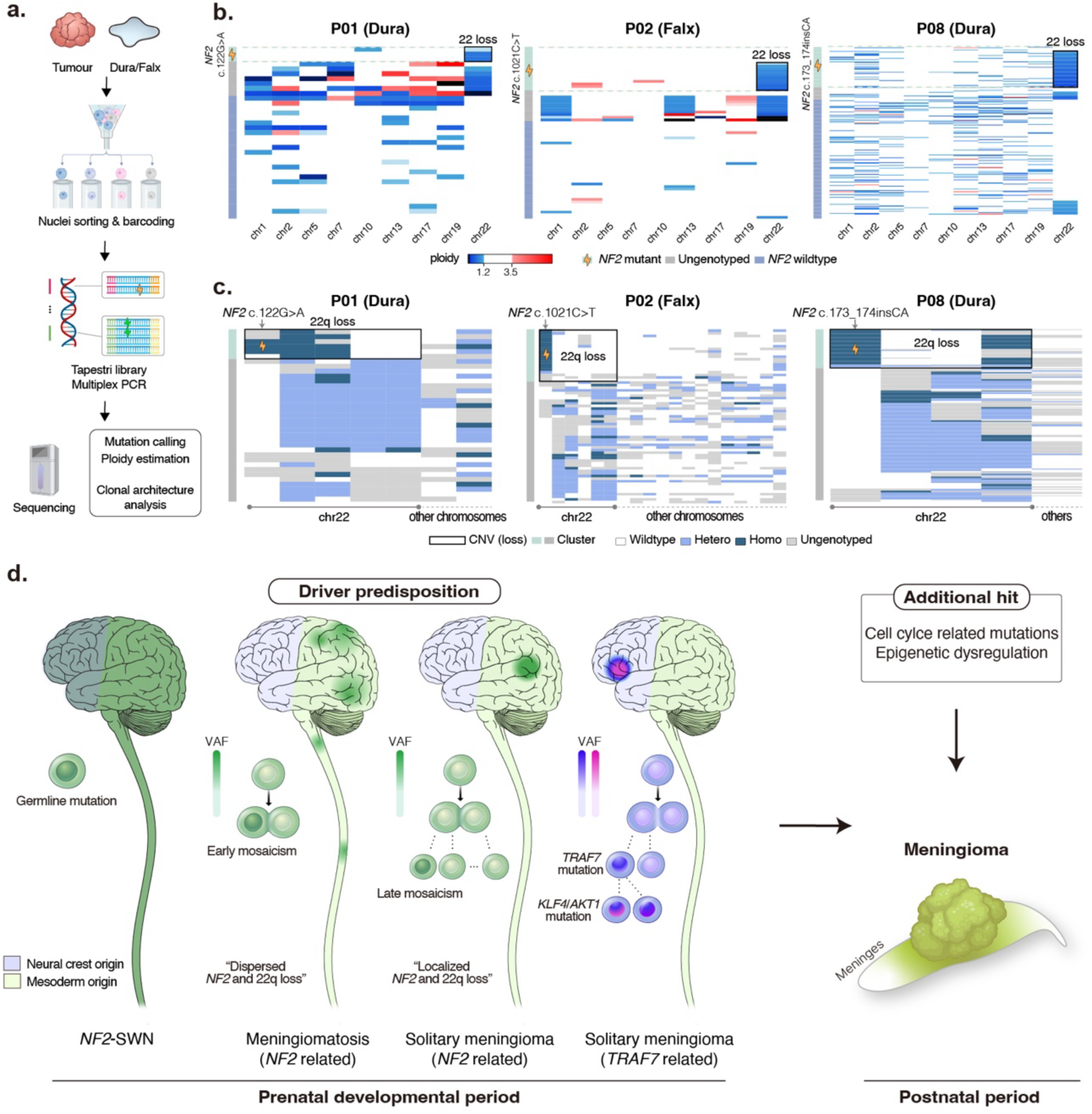
A framework for the developmental origin and spatial patterning of human meningioma. **a,** Workflow for single-nucleus DNA sequencing of paired tumour and tumour-free meninges using the Tapestri droplet-based platform. **b, c,** Estimated ploidy heatmaps (**b**) and genotype distributions (**c**) of *NF2* mutation status and 22q copy-number state in nuclei isolated from normal meninges across three samples, showing co-occurrence of *NF2* mutation and 22q loss in 8.6–20.6% of profiled nuclei. **d,** Conceptual model illustrating how the timing and spatial extent of developmental meningeal mosaicism establish pre-neoplastic fields of varying dimensions, shaping tumour multiplicity and anatomical distribution, with subsequent postnatal alterations driving clinical tumour manifestation.

Among 35, 67 and 247 nuclei sequenced, co-occurrence of *NF2* mutations and 22q loss inferred by read-depth estimation was observed in 3 (8.6%), 10 (14.9%), and 51 (20.6%) of cells, respectively (**Fig. 5b**). Based on genotype-driven cell clustering, *NF2* driver mutations co-occurred in 42.9% (3/7), 81.0% (17/21), and 96.5% (55/57) of clusters exhibiting 22q loss, respectively (**Fig. 5c**). Considering the inherent genotyping errors and allelic dropout associated with single-nucleus DNA sequencing platforms, these findings suggest that *NF2* mutation and 22q loss may co-occur early during meningeal development.

Despite evidence for early acquisition of major driver events, meningiomas typically manifest clinically decades later, suggesting that additional postnatal alterations may contribute to tumour progression. To explore this possibility, we performed a network-based functional analysis focusing on tumour-private clonal mutations that were not shared with the adjacent meninges (**Extended Data Fig. 8a**). Gene ontology analysis (**Methods**) revealed enrichment in biological processes related to cell cycle regulation (‘Positive regulation of cell cycle phase transition’ (adjusted *P* = 0.02) and ‘Hippo signaling regulation pathways’ (adjusted *P* = 0.02)) and epigenetic dysregulation (‘Histone H3 acetylation’ (adjusted *P* = 0.03) and ‘Positive regulation of double-strand break repair’ (adjusted *P* = 0.03)) (**Extended Data Fig. 8b**). These pathways may therefore represent candidate postnatal events contributing to tumour progression beyond the initial developmental hits.

### Expanding to NF2-associated syndromic disease

The spatiotemporally expanded role of *NF2* mutations in meningioma suggests a biological link between solitary meningioma and *NF2*-related syndromic disorders, including *NF2*-SWN and MM. In addition to inherited or *de novo* germline *NF2* mutations, recent studies estimate that mosaic *NF2* mutations detectable in peripheral blood account for approximately 25–60% of cases (**Extended Data Fig. 9**)^12^. These observations support substantial biological overlap, with mosaic *NF2* mutations potentially conferring genetic and histopathological features shared between solitary meningioma and syndromic disease. Indeed, solitary and multiple meningiomas showed no significant differences in driver mutation spectra (**Extended Data Fig. 10a)**, number of subclonal mutations (6.7 ± 2.9 vs 6.2 ± 3.8, *P* = 0.45, **Extended Data Fig. 10b**), mutational signatures (cosine similarity > 0.99, **Extended Data Fig. 10c**), copy-number alteration profiles, or histological subtypes. Instead, our data support a model in which the timing of driver mutation acquisition and its anatomical distribution contribute to tumour multiplicity and spatial extent.

Accordingly, the distinction between *NF2*-SWN and MM may be more appropriately interpreted as a continuum influenced by the timing and extent of *NF2* mosaicism, rather than as strictly discrete entities defined solely by clinical features. For example, patients P01 and P04, who presented with multiple meningiomas along the craniospinal axis without schwannoma, are currently classified as MM; however, subsequent development of schwannomas could potentially result in reclassification as *NF2*-SWN. Together, these findings suggest that the developmental timing and anatomical distribution of *NF2* mosaicism contribute to the clinical spectrum and spatial extent of NF2-associated disease.

## Discussion

In this study, we provide direct genomic evidence that developmental somatic mosaicism underlies a substantial proportion of human meningiomas, based on matched analyses of tumour and tumour-free meningeal tissues. Approximately 82% of meningioma patients harboured shared low-VAF driver mutations in tumour-free meninges, consistent with developmental mosaicism, accounting for 88% and 75% of cases with *NF2-* and *TRAF7-*driven tumours, respectively. These shared low-VAF driver mutations were detectable in tumour-free meninges and supported by orthogonal validation approaches including ultra-deep amplicon sequencing, phylogenetic reconstruction, and single-cell analyses. Our results suggest that clinically distinct disease entities, including *NF2*-SWN, MM, and solitary meningioma, share overlapping developmental genetic origins and may therefore be interpreted within a unified developmental framework.

Previous mouse studies have established that biallelic *Nf2* inactivation in primordial meningeal progenitor cells (but not in adult meningeal cells) is both necessary and sufficient for meningioma formation, defining a restricted embryonic window permissive for tumour initiation^22^. Our findings provide the first direct genomic evidence that an analogous developmental mechanism operates in human meningiomas. In particular, the coexistence of *NF2* mutations and 22q loss within tumour-free meninges indicates that biallelic *NF2* inactivation can arise before overt tumour formation. However, considering the long latency between development and clinical presentation, these early events are unlikely to be sufficient for full tumourigenesis. Consistent with epidemiological models proposing multiple rate-limiting steps in meningioma development^21^, our data support additional lineage-restricted post-developmental events during tumour progression. Candidate mechanisms identified in our analyses include SBS15-associated mutagenesis, cell-cycle dysregulation, and epigenetic alterations (**Extended Data Fig. 8**), although their precise mechanistic contributions remain to be established.

A developmental perspective provides a biologically grounded framework for understanding the *NF2*-associated disease spectrum (**Extended Data Fig. 9**). Current diagnostic systems, including the Manchester criteria^12^, rely heavily on phenotype, family history, and blood-based testing, but do not fully account for lineage-specific developmental mechanisms. Our findings suggest that the timing and anatomical distribution of mosaic *NF2* mutations substantially influence tumour multiplicity and disease extent. Within this framework, mosaic mutations arising before versus after developmental lineage divergence would be expected to generate distinct clinical trajectories and oncological burdens. More broadly, *NF2*-related disorders may be conceptualized along a continuous mutational spectrum encompassing germline, de novo, early mosaic, late mosaic, and postnatal somatic variants, although the precise diagnostic boundaries will require prospective genetic–clinical correlation. We therefore anticipate that integrating developmental mosaicism into diagnostic classification may facilitate a more biologically coherent interpretation of *NF2*-related disease.

While developmental mosaicism has previously been implicated in rare congenital disorders and paediatric tumours^23–25^, our study provides direct human genomic evidence supporting the concept that developmental mosaicism can also contribute to adult-onset neoplasia. To date, inherited germline predisposition has explained only a minority of patients across many common tumour types, including approximately 5–10% of breast cancers associated with *BRCA*1/2 mutations and ∼1% of colorectal cancers linked to germline *APC*-associated familial adenomatous polyposis^26^. Our findings raise the possibility that developmental mosaic mutations may represent an additional and previously underappreciated layer of cancer predisposition that is not captured by conventional germline testing. Developmental mosaicism may therefore represent a cryptic intermediate state between germline predisposition and purely postnatal somatic tumourigenesis in that it confers cancer predisposition only within a spatially restricted region. This concept of ’field cancerization’ has been raised by recent deep sequencing studies of normal tissues including the endometrium, oesophagus, and colorectal epithelium, yet actual progression to tumour formation has not been clearly established^27–29^. Our data provide direct evidence that developmental driver mutations persist in normal meninges to the adjacent tumours they give rise to. We anticipate that developmental mosaicism may similarly underlie other adult neoplasms arising from anatomically and embryologically constrained tissues.

Several limitations remain. First, although our findings strongly support developmental mosaicism within meningeal tissues, the contribution of arachnoid tissue itself could not be directly assessed because sufficient material was difficult to obtain surgically. Second, the number of higher-grade meningiomas in our cohort was limited, and additional studies will be required to determine whether similar developmental mechanisms contribute to more aggressive disease states. Finally, the mechanistic basis underlying preferential localization of *TRAF7*-, *KLF4*-, and *AKT1*-mutant tumours within specific meningeal compartments remains unresolved.

In conclusion, our study suggests that many human meningiomas acquire tumour-initiating mutations during development and demonstrates that ultra-low-frequency mosaic mutations can persist in tumour-free meninges at VAFs as low as 0.02%. Collectively, these findings position developmental somatic mosaicism as a previously underappreciated mechanism underlying human meningioma and potentially contributing to adult oncogenesis more broadly.

## Data Availability

Raw sequencing data generated in this study, including WES, AS, WES of single-cell-cloning, Tapestri single-nucleus DNA sequencing, and PacBio long-read sequencing, have been deposited at the NCBI SRA (PRJNA1469910) and KRA (KAP242312). Processed Tapestri single-nucleus DNA sequencing data in .h5 format are available at FigShare (https://doi.org/10.6084/m9.figshare.32386740).

## Code Availability

Code for the data processing and analysis is provided at the author’s GitHub repository (https://github.com/goldpm1/meningioma)

## Acknowledgement

We thank MID (Medical Illustration & Design), part of the Medical Research Support Services of Yonsei University College of Medicine, for excellent support with medical illustration. We also thank InDNA, JSLink Inc., TheragenBio, Corp., and Macrogen, Inc. for their contributions to library preparation, sequencing, and sequencing data generation. Figure 4d and 5a were created with BioRender.com.

## Author Contributions

S.-G.K., Y.C. and S.K. conceived and designed the study. S.H.K. and T.C. performed the microscopic investigation and pathological assessment. J.H.M., S.C., T.-H.R., E.H.K., C.K.L., D.A.S., S.Y., Y.H., K.N.K., J.H.C. and S.-G.K. collected the samples. Y.C., T.K. and S.K. performed the bioinformatic analyses and interpreted the data. Y.C. and H.J.C. designed and performed the experimental verification. Y.C. and H.J.C. drafted the manuscript. S.K. and S.-G.K. critically revised the manuscript. All authors reviewed the submitted version of the manuscript. S.-G.K. and S.K. approved the final version on behalf of all authors.

## Methods

### Patient samples and clinical data

Matched pairs of tumours, adjacent meninges, and blood were collected from 27 individuals who underwent surgical resection for intracranial or spinal meningiomas at Severance Hospital (Seoul, Republic of Korea). This cohort comprised 22 patients with histologically confirmed tumour-free adjacent meninges and 5 patients with microscopically tumour-infiltrated meninges (C01–05). Surgical resection achieved Simpson grade 1 (convexity) or 2 (ventricle, skull base, and spinal canal)^30^ to ensure the total removal of the tumour. Meningeal samples were collected from the site furthest from the tumour within the surgical window. To prevent cerebrospinal fluid leakage, the collection was limited to a single region. Additionally, tumour-free samples from the cerebral cortices (n=2) and lateral ventricular walls (n=2) were included. More than two independent neuropathologists determined the histologic diagnosis and grading following the 2021 WHO Classification of Tumours of the CNS. Determination of whether tumour-free or not was made by extensive inspection through a 100–200x high-power field (HPF)^31^.

For P01, five different tumours from convexity, parasagittal, and falx were collected. These were designated T1–5, each of which represents the frontal falx tumour, parietal falx tumour, parietal convexity tumour, posterior parasagittal, and posterior tumour, respectively. For P02, four intracranial tumours—comprising two falx meningiomas and two relatively small parasagittal meningiomas—were initially collected and designated T1–T4. One year later, an additional tuberculum sella meningioma (T5) was resected via an endoscopic transsphenoidal approach (TSA). For P03, samples were obtained from three separate surgical procedures, yielding one convexity meningioma and two thoracic spinal meningiomas. A small vestibular schwannoma (VS) in P03 was managed conservatively and not surgically resected.

We gathered clinical data through electronic medical record (EMR), including age, sex, and history of previous operations for neuro-oncologic disease with appropriate de-identification. All the locations where samples were collected were noted in the EMR at the time of surgical resection. The study was approved by the Institutional Review Board of Severance Hospital (IRB No. 4-2021-1319) and conducted following the Declaration of Helsinki. All patients provided written informed consent.

### Whole-exome sequencing

Genomic DNA (1.0 µg) was extracted from samples, and paired-end sequencing libraries were generated using the Agilent SureSelect Target Enrichment protocol (Agilent Technologies, Santa Clara, CA, USA). The SureSelect Human All Exon V8 probe set was utilized in all cases. DNA quantity and quality were assessed using PicoGreen and agarose gel electrophoresis. Genomic DNA was diluted in EB Buffer and sheared to a target peak size of 150–200 bp using a Covaris LE220 focused-ultrasonicator (Covaris, Woburn, MA, USA) with the following settings: frequency sweeping mode, 10% duty cycle, intensity 5, 200 cycles per burst, and a duration of 60 seconds for 6 cycles at 4°C–7°C. Fragmented DNA underwent end-repair, 3′-adenylation, and ligation with Agilent adapters. Following ligation assessment, products were PCR-amplified. For exome capture, 250 ng of the DNA library was hybridized with the SureSelect all exon capture baits at 65°C for 24 hours. Captured DNA was washed, amplified, and quantified by qPCR using the KAPA Library Quantification Kit (Roche, Basel, Switzerland). Library quality was validated using the TapeStation DNA ScreenTape D1000 (Agilent). Sequencing was performed on the Illumina NovaSeq X platform (Illumina, San Diego, CA, USA).

### Variant calling

Sequence reads were aligned to the hg38 reference genome using BWA-MEM (v.0.7.17). Duplicate reads were marked and removed using GATK (v.4.2.3.0) MarkDuplicates. Base quality score recalibration (BQSR) and indel realignment were performed using GATK BaseRecalibrator and LeftAlignIndels, respectively. Somatic variants were called using GATK Mutect2 in tumor-normal mode with matched peripheral blood. Initial calls were strictly filtered using GATK FilterMutectCalls (settings: --min-median-base-quality 20, --min-median-mapping-quality 20, --min-reads-per-strand 1, and --max-events-in-region 1). Variants were further excluded if they had a read depth < 30, were multiallelic, or were located in repeat regions. For non-tumour samples, we investigated the presence of alternate reads at candidate sites using pysam (v.0.22.0). Variants matching those in the corresponding tumour were rescued if they were monoallelic and met a base quality threshold > 20 (**Extended Data Fig. 5**).

### Ultra high-depth amplicon sequencing

For the 22 dura, 2 cortex, and 2 ventricular wall samples, ultra-high-depth (∼1,000,000x) massive parallel amplicon sequencing (MPAS) was performed for 22 candidate driver mutation sites. Additionally, single-site amplicon sequencing (∼10,000,000x) was conducted for five specific positions—*TRAF7* (chr16:2,175,609 and chr16:2,176,145), *KLF4* (chr9:107,487,067), *AKT1* (chr14:104,780,214), and *NF2* (chr22:29,673,365)—where MPAS coverage was suboptimal. Targeted regions were amplified using PrimeSTAR GXL high-fidelity DNA polymerase (Takara Bio, Kusatsu, Japan). Sequencing libraries were constructed using the TruSeq Nano DNA High Throughput Library Prep Kit (Illumina). Briefly, 100 ng of amplicon DNA underwent end-repair, ‘A’-tailing, and ligation with TruSeq DNA UD Indexing adapters. Purified and PCR-enriched libraries were quantified via qPCR and validated on a 4200 TapeStation (Agilent). Final sequencing was performed on the Illumina NovaSeq X system.

### Variant detection in amplicon sequencing

Raw reads from MPAS and single-site amplicon sequencing were aligned to the GRCh38 reference genome using BWA-MEM (v.0.7.17). Base quality scores were recalibrated using GATK BaseRecalibrator (v.4.2.3.0). Duplicate reads were retained to preserve the quantitative nature of the amplicon data. Pysam (v.0.22.0) was used to analyze base compositions at target positions. Reads were excluded from analysis if they met any of the following criteria: i) presence of hard or soft clipping at the read ends, ii) variant location within 3 bp of either end of the read, iii) base quality < 20 at the variant site, or iv) presence of clustered variants within a 3 bp window. Strand bias was determined if one strand represented less than 5% of the other strand, allowing enough tolerance to account for amplification bias during ultra-high-depth sequencing.

To distinguish true positives from sequencing artifacts, we employed a binomial statistical model for each candidate variant. We assumed that the number of false-positive reads at a given site follows a binomial distribution:

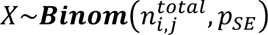

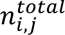 and *VAF_i,j_* stands for total read count and variant allele frequency for variant *j* of sample *i*, respectively. *p_SE_* denotes the background sequencing error rate, estimated as the median of *VAF_i,j_* for each sample *i*.

The statistical significance of each observed variant was determined based on its cumulative probability within the null distribution *X*. Variants falling within the top 5th percentile (*P* ≥ 0.95) were classified as true positives. Variants within the top 5th to 10th percentile (0.95 > *P* ≥ 0.90) were categorized as indeterminate (represented in grey in the heatmaps). Those falling within the remaining 90% of the distribution (0.90 > *P*) were regarded as sequencing artifacts. Visualization, including heatmaps and swarm plots, was performed using the Seaborn (v.0.12.2) and Matplotlib (v.3.7.1) packages in Python.

### Long-read sequencing

For patient P12, who harboured 22q loss without an identifiable NF2 mutation, high-molecular-weight (HMW) genomic DNA (gDNA) was extracted using the Promega Wizard HMW DNA Extraction Kit (Promega, Madison, WI, USA), Nanobind panDNA Kit (Pacific Biosciences, CA, USA), or NEB Monarch HMW DNA Extraction Kit (New England Biolabs, Ipswich, MA, USA), depending on the sample type. To minimize mechanical shearing during lysis and purification, wide-bore tips and gentle inversion were utilized throughout the protocol. DNA concentration was determined using the Qubit dsDNA HS Assay Kit (Thermo Fisher Scientific, MA, USA), and size distribution was assessed on an Agilent 2100 Bioanalyzer with the Genomic DNA ScreenTape (Agilent Technologies, CA, USA) to ensure fragments exceeded 48 kb. For library preparation, gDNA was sheared to an average size of 15–20 kb using a Covaris g-TUBE device (Covaris, MA, USA) and purified with SMRTbell Clean-up Beads (Pacific Biosciences, CA, USA). SMRTbell libraries were constructed using the SMRTbell Prep Kit 3.0 (Pacific Biosciences, CA, USA) following the manufacturer’s instructions and sequenced on the PacBio Revio system to generate high-fidelity (HiFi) reads (Q20+). 5-Methylcytosine (5mC) at CpG sites was predicted from the HiFi reads using pbjasmine (v.2.2.1), and unaligned BAM files containing methylation data were demultiplexed using lima (v.2.10.0).

Additionally, total RNA was isolated from tumour tissue of patient P12, who harboured 22q loss without an identifiable NF2 mutation, using the Hybrid-R kit (GeneAll, Seoul, Republic of Korea) according to the manufacturer’s instructions. As controls, tumour tissues from two NF2-wildtype meningiomas, one microcystic and one meningothelial, were processed in parallel; these control cases were not included in the main study cohort. Tissue samples were lysed with RiboEx solution, and pure RNA was recovered via glass-fiber membrane column purification. RNA concentration and purity were evaluated using a NanoDrop spectrophotometer and a Qubit 4 Fluorometer with the RNA HS Assay Kit (Thermo Fisher Scientific, MA, USA), while RNA integrity was confirmed on an Agilent 2100 Bioanalyzer using the RNA 6000 Nano Kit (Agilent Technologies, CA, USA). For long-read transcriptome sequencing, Kinnex Iso-Seq libraries were prepared using the Iso-Seq Express 2.0 Kit (Pacific Biosciences, CA, USA). Briefly, full-length cDNA was synthesized with specific handles, and 5’ barcodes were added during cDNA amplification. To optimize sequencing throughput, Kinnex adapter ligation was performed to concatenate cDNA arrays. Final library quantification and quality assessment were performed using the Qubit dsDNA HS Assay Kit and the Agilent 2100 Bioanalyzer with the DNA 12000 Kit, respectively. Libraries were sequenced on a PacBio Revio system (Pacific Biosciences, CA, USA).

### Detection of structural variation using long-read sequencing

PacBio HiFi reads and Iso-Seq data were aligned to the T2T-CHM13 reference genome using pbmm2 (v.1.13.1). Alignment parameters were optimized using the --preset CCS for genomic DNA and --preset ISOSEQ for RNA data. Genomic structural variations (SVs) were identified using Sniffles2 (v.2.2). Transcriptomic data were processed following the PacBio Iso-Seq pipeline for isoform classification and downstream characterization. Candidate genomic rearrangements and structural alterations involving *NF2* were manually inspected and visualized using the Integrative Genomics Viewer (IGV).

### Allele-specific copy number profiling

Allele-specific somatic copy number alterations (SCNAs) were detected using cnv_facets (v.0.16.0), with matched peripheral blood samples serving as normal controls. Tumour purity and ploidy estimates were further validated using the Sequenza R package (v.3.0.0). The resulting binning data and read-depth log-fold changes were utilized for downstream clonality analysis. Following the molecular classification criteria proposed by Bayley et al.^5^, tumours were categorized into three subgroups: MenG A (neither 1p nor 22q loss), MenG B (22q loss only), and MenG C (combined 1p and 22q loss). Sample-wise arm-level copy number changes were visualized using the ComplexHeatmap R package (v.2.14.0).

### Assessment of tumour contamination in bulk datasets

To evaluate potential tumour cell contamination within the normal tissue samples, quantitative profiling for all patients harboring shared driver mutations, including tumour infiltrated cohort (C01–05). The number, VAF, and proportions of shared non-driver mutations between the two cohort was evaluated. Statistical significance of the differences in these quantitative metrics between the two groups was evaluated using the non-parametric Mann–Whitney U test. Additionally, *in silico* Contamination levels were assessed using TINC (v.0.1.0)^15^ and MeTel (v.1.0)^16^. For TINC analysis, variant call format (VCF) files and copy number profiles estimated by Sequenza were used as inputs to calculate Tumour-in-Normal (TIN) scores. The resulting TIN scores were compared across the cohort and visualized using the Matplotlib (v.3.7.1) library in Python. Additionally, MeTel was employed for pairwise comparisons between tumours and matched meningeal samples. The MeTel model was specifically tailored for meningioma by designating *NF2*, *TRAF7*, *KLF4*, and *AKT1* as key driver genes, with prior allele frequencies integrated from gnomAD v4.1. The classification scores for all sample pairs were presented as boxplots using Matplotlib.

### Clonality analysis

For each patient, somatic variant data were obtained from WES of matched tumour and meningeal samples. Allele-specific copy-number profiling and tumour purity estimates were derived using cnv_facets. Clonal architecture was subsequently analysed using PyClone-vi, employing the beta-binomial model with all other parameters set to default. Clonal relationships were visualised via two-dimensional scatter plots, where the variant allele frequencies (VAFs) of the meninges and tumour were plotted on the x- and y-axes, respectively. For somatic variants of exceptionally low frequency that fell below the detection threshold of standard whole-exome sequencing (WES)—specifically encompassing eight low-level driver mutations identified across seven patients—high-depth targeted amplicon sequencing data were utilised. Each cluster was assigned a distinct colour, with circles representing shared variants and triangles representing non-shared variants. Clusters exhibiting the highest cellular prevalence within the tumour were classified as clonal, while all remaining clusters were designated as subclonal. Based on these definitions, the proportion of shared clonal non-driver mutations (calculated as the ratio of tumour-clonal non-driver mutations shared with the meninges) was utilised for subsequent statistical comparisons.

### Mutational signature analysis

For 22 individuals, somatic variants were categorised into three distinct groups: clonal mutations shared with the meninges, clonal mutations unique to the tumour, and subclonal mutations unique to the tumour. The mutational context matrix was constructed using SigProfilerMatrixGenerator (v.1.2.0). Mutational signature analysis was subsequently performed using MuSiCal (v.1.0.0)^32^, an NMF-based deconvolution framework. Following de novo signature discovery, variants were assigned to established COSMIC signatures (v.3.3). The resulting data were visualised as stacked bar plots using the Matplotlib (v.3.7.1) library in Python.

### Single-cell expansion and DNA sequencing

From two individuals (P01 and P21), each representing *NF2* (+) and *KLF4*(+), *TRAF7*(+) meningioma, bulk tumour spheres were successfully constructed using a previously reported method for tumour sphere isolation from the human brain^33^. Out of five tumours of P01, we selected T5 (posterior tumour) because of its highest viability. Within hours after the operation, meningioma-derived tumour spheres (TS) were generated from the patient’s tumour tissue. The meningioma TSs were maintained in complete media consisting of Dulbecco’s Modified Eagle’s Medium/F12 (Mediatech), 1x B27 (Invitrogen), 20 ng/mL basic fibroblast growth factor, and 20 ng/mL epidermal growth factor (Sigma-Aldrich). To establish separate clonal lines, meningioma TS were dissociated into single cells using Accutase. The cells were then diluted at a ratio of 1:100 and plated into 96-well plates. After 3 days, wells containing single cells were manually selected under a microscope, and the cells were gradually expanded.

Following the quality and quantity check using the cell counter, we extracted genomic DNA from each clone with the QIAamp micro-DNA kit (Qiagen) following the manufacturers’ instructions. For 8 clones of P01, we established an exome library without further genomic amplification. For 8 clones derived from P21, we exploited primary template-directed amplification (PTA), due to the lack of the DNA amount to execute conventional WES. PTA was performed using the ResolveDNA™ Whole Genome Amplification Kit (BioSkryb PN100136, NC, USA) following the manufacturer’s protocol. All PTA reagents were added step-by-step according to the provided instructions. The lysed samples, along with the enzymes and terminators, were incubated on a thermal cycler at 30°C for 10 hours. This was followed by enzyme deactivation at 65°C for 3 minutes. Finally, bead purification of the amplicons was conducted as instructed by the manufacturer. Then, ∼300x exome libraries (Agilent SureSelect XT HS) were prepared for all clones of two individuals. The method of exome capture and sequencing is identical to the aforementioned. FASTQ files were processed and read alignment to GRCh38 was conducted accordingly.

For each individual, we collected ∼900x bulk meninges and ∼300x clonally expanded tumour cells. GATK v4.2.3.0 HaplotypeCaller with GVCF and Ploidy 2 options was conducted for somatic mutation calling. Combined genotyping for meninges and single-cell derived tumour samples was executed through GATK GenomicDBImport and GenotypeGVCFs, followed by ApplyVQSR for SNP and indel calls, separately. To increase confidence, we additionally applied the filtering process by binomial distribution model. In each pre-genotyped result, we investigated the cumulative distribution function as follows:

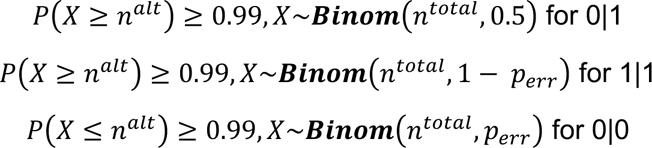

where*n^alt^* and *n^total^* denotes alternate and total read counts.

Then, we selected the variants where the genotype of the tumour clone is 0|1 or 1|1 and the genotype of the dura is 0|0. Variants shared in all tumour clones were nominated as tumour clonal mutations. Data visualization as a heatmap was generated with the seaborn package.

### Phylogenetic tree reconstruction

Phylogenetic relationships among clones within the same individual were reconstructed based on a matrix of total depth and variant-supporting alternate reads. Tree inference was performed using the Sequoia R script, which employs MPBoot for maximum parsimony analysis. Since germline variants had been removed during the preceding analytical stages, the --germline_cutoff parameter was set to its minimum value to prevent the inadvertent loss of somatic variants. For variants identified in bulk meningeal samples, the alternate read count was adjusted to 50% of the total depth. This adjustment was made to comply with the expected heterozygous state in single-cell sequencing data. Phylogenetic trees were initially generated in Newick format using the ggtree R package (v.3.6.2) and were subsequently finalised manually as diagonal cladograms.

### Droplet-based single-nuclei DNA sequencing

To perform high-throughput single-cell DNA sequencing using the Tapestri platform (MissionBio, San Francisco, CA, USA), single nuclei were isolated from cryopreserved tumour and meningeal tissue samples from patients P01, P02 and P08. Tissues (50–100 mg) were minced on ice and homogenised in 1 mL of lysis buffer (comprising 0.2% Triton X-100, 1 × Roche protease inhibitor, 1 mM DTT, and 0.2 U/µL RNAsin in 2% BSA in PBS). The nuclei pellet, recovered via centrifugation at 500g for 5 minutes at 4°C, was resuspended in 400 µL of suspension buffer (1 mM EDTA, 0.2 U/µL RNAsin, and 2% BSA in PBS). After filtration through a 40µm strainer and DAPI staining (0.5 µg/mL), DAPI-positive nuclei were isolated using a flow cytometer (BD Biosciences, Franklin Lakes, NJ, USA) following a secondary 20µm filtration step.

A custom multiplex amplicon panel was designed via Tapestri Designer (MissionBio), incorporating exons of *NF2*, *TRAF7*, *KLF4*, and *AKT1*, alongside target regions on chromosome 22q, into the standard Tapestri glioma panel. This final configuration comprised 297 amplicons and 1,272 targets (total size 52.9 kb). Approximately 100,000 nuclei per sample were loaded onto a Tapestri microfluidic cartridge for encapsulation with lysis reagents, followed by protease inactivation. Within the droplets, cell-identifying molecular barcodes were attached to targeted genomic regions using gene-specific primers and hydrogel beads. After UV-mediated primer cleavage and targeted PCR amplification, the emulsions were broken and the DNA was purified using AMPure XP beads (Beckman Coulter, Brea, CA, USA). Following the addition of Illumina adapter sequences via a 10-cycle PCR and a final purification, library quality was verified on a 4200 TapeStation (Agilent Technologies, Santa Clara, CA, USA). Sequencing was performed on an Illumina NovaSeq 6000 system (Illumina, San Diego, CA, USA) with 2×150 bp paired-end reads. Experimental procedures from library preparation to sequencing were conducted by JSLink, Inc. (Seoul, Republic of Korea).

### Integrated genotyping and ploidy inference at the single-nucleus level

The Tapestri bioinformatics pipeline (v.2) was utilised for adapter trimming, alignment, and barcode assignment to individual nuclei. To distinguish these droplet-based data from whole-genome sequenced clones, a specific genotyping and ploidy inference framework was applied using the Mosaic software (v.3.7). Single-nucleus genotyping was performed using a GATK-based algorithm, and strict quality filters were applied to ensure the integrity of the results. Specifically, nuclei were excluded if they exhibited a read depth < 10 or a genotype quality (GQ) score < 30. Furthermore, we filtered out genes genotyped in fewer than 30% of total cells and those exhibiting alternate alleles in fewer than 1% of the population.

For the analysis of chromosomal alterations, copy-number states were inferred for each nucleus using the filter_amplicons command (completeness = 50, read depth = 10). Following the normalisation of read depth across amplicons and individual nuclei, ploidy levels were estimated via the cnv.compute_ploidy command, with wild-type nuclei utilised as diploid controls. Heatmaps integrating estimated ploidy with NF2 genotypes were generated, providing a high-resolution map to reconstruct the temporal order of mutational events and loss of heterozygosity (LOH) at the single-nucleus level.

### Network analysis for clonal mutations

To investigate the molecular function of clonal variants other than *NF2*, *TRAF7*, *AKT1*, and *KLF4*, we performed a gene enrichment analysis for GO biological process, Wiki, Reactome, and KEGG pathway using ClueGO (v.2.5.10) in Cytoscape (v.3.10.1). Clonal, nonshared variants derived from the previous clonality analysis of WES data were utilized. We set parameters as Evidence All_without_IEA (inferred from electronic annotation), GO tree interval ranging from 3 to 8, and at least 5 genes, 4% of genes for GO term selection and a kappa score of 0.4. We adjusted the P–value using the Bonferroni step-down with a significance criterion of 0.05.

### Statistical testing

The statistical testing in **Fig. 2** and **Extended Data Fig. 6** (Mann Whitney test and Pearson’s correlation test) was evaluated through the scipy package. Statistical significance is shown by asterisks (\**P* ≤ 0.05, \*\**P* ≤ 0.01) in the figure legends. Correlations between dura and tumour VAFs were computed per patient using Pearson correlation. Fisher’s z-transformation was applied and 95% CIs were calculated as 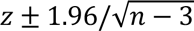. Pooled correlations were estimated using fixed-effect meta-analysis of Fisher z-transformed coefficients with inverse-variance weighting (*w* = *n* − 3), then back-transformed to r values. One-sided tests were used to evaluate whether pooled correlations were greater than zero.

## Extended Data

**Extended Data Fig. 1.**
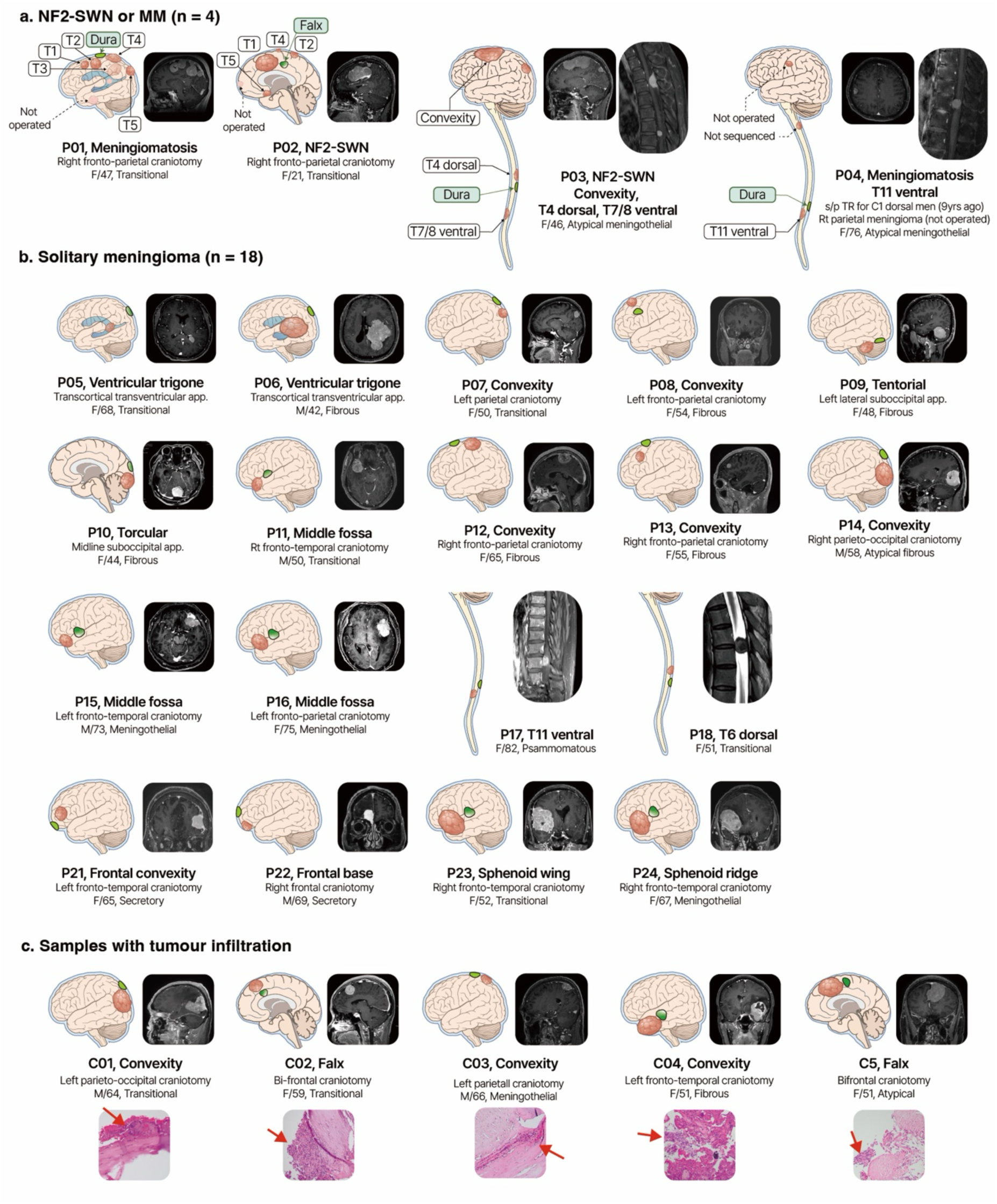
Patient cohort and tissue collection. The patient cohort and the location of tissue collection, including tumour and dura, are shown. A total of 22 individuals with 32 tumours are included; four of these are anteriorly located meningiomas harbouring *TRAF7* mutations, whereas *NF2* variants were identified in the remaining cases. Five individuals with tumour-infiltrated meninges served as positive controls (PC). Clinical diagnosis, surgical approach, and the history of prior surgery are shown for each individual. **a,** *NF2*-SWN or MM (n = 4), **b,** solitary meningioma (n = 18), and **c,** individuals with pathologically confirmed tumour infiltration used as positive controls (PC; n = 5). Arrows indicate regions of tumour infiltration on histology.

**Extended Data Fig. 2.**
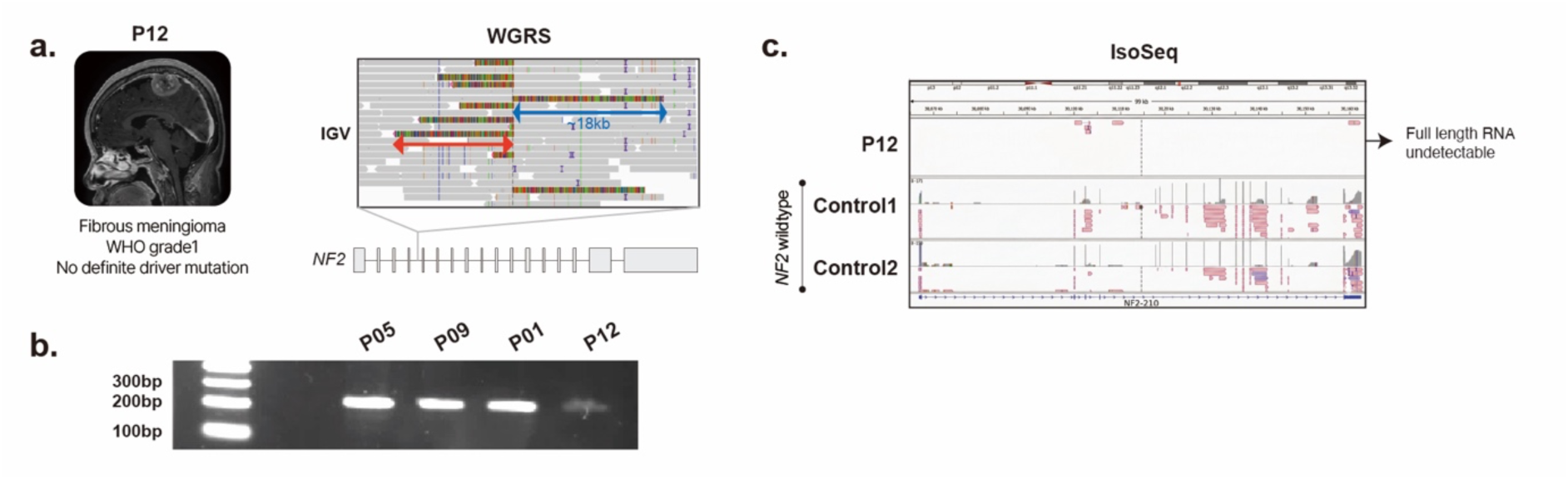
Complex structural variant encompassing *NF2*. Long-read sequencing was performed on patient P12, who harboured 22q loss without an identifiable *NF2* driver mutation, to investigate potential hidden genomic alterations. **a,** IGV view showing bidirectional soft-clipped reads (∼18 kb and ∼14 kb) within intron 4 of *NF2*, with breakpoints separated by 140 base pairs. **b,** Gel electrophoresis of a PCR product spanning the breakpoint, indicating a partial genomic disruption in P12 that was absent in control samples. **c,** PacBio Iso-Seq analysis of tumour RNA showing markedly reduced *NF2* expression in P12 relative to two *NF2*-wildtype control tumours. Notably, no full-length *NF2* transcript isoform was detected in P12. **IGV**; Integrated Genome Viewer

**Extended Data Fig. 3.**
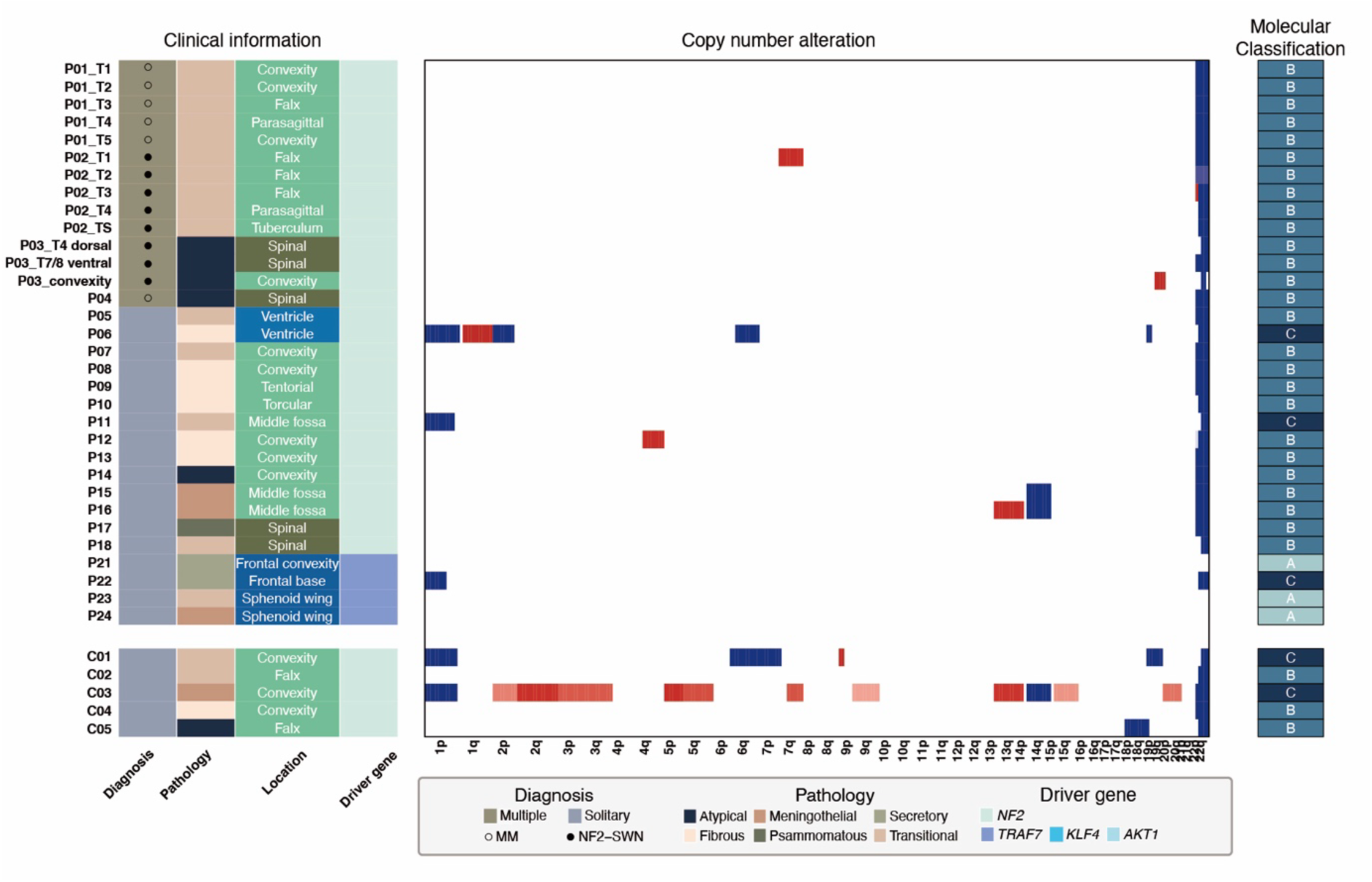
Copy number alteration of tumours and meninges. For each sample, clinical information and genomic features, including driver alterations and copy-number profiles, are shown. The colour bar indicates the amplitude of copy number relative to the diploid baseline, with red and blue representing gains and losses, respectively. Molecular classification from Bayley *et al.*^5^ is provided in the right hand column.

**Extended Data Fig. 4.**
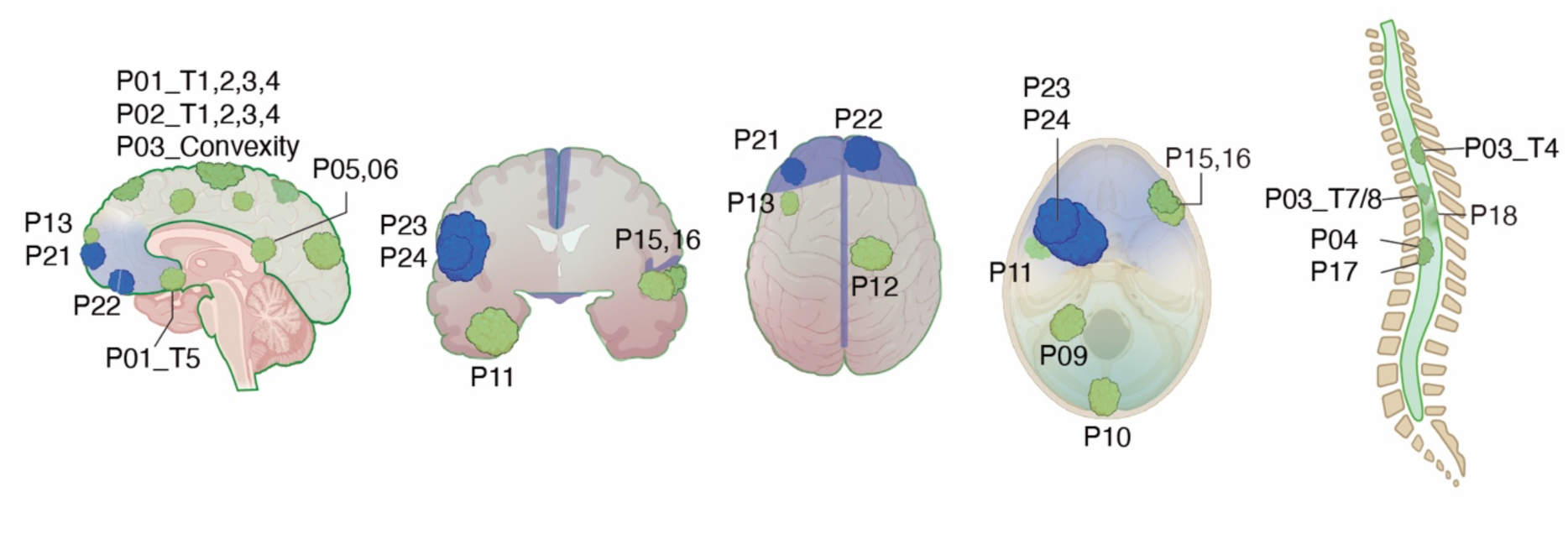
Anatomical preference of meningioma subgroups defined by driver mutations. Four non-*NF2*-related meningiomas were located in the anterior cranial vault and anterior skull base, regions often considered neural crest derivatives (blue-shaded regions). In contrast, *NF2*-related meningiomas were distributed throughout the entire craniospinal axis (green-shaded regions). Tumour locations are labelled by case ID; *NF2*-related tumours are marked in green, whereas non-*NF2* tumours (*TRAF7* with *KLF4* or *AKT1*) are marked in blue.

**Extended Data Fig. 5.**
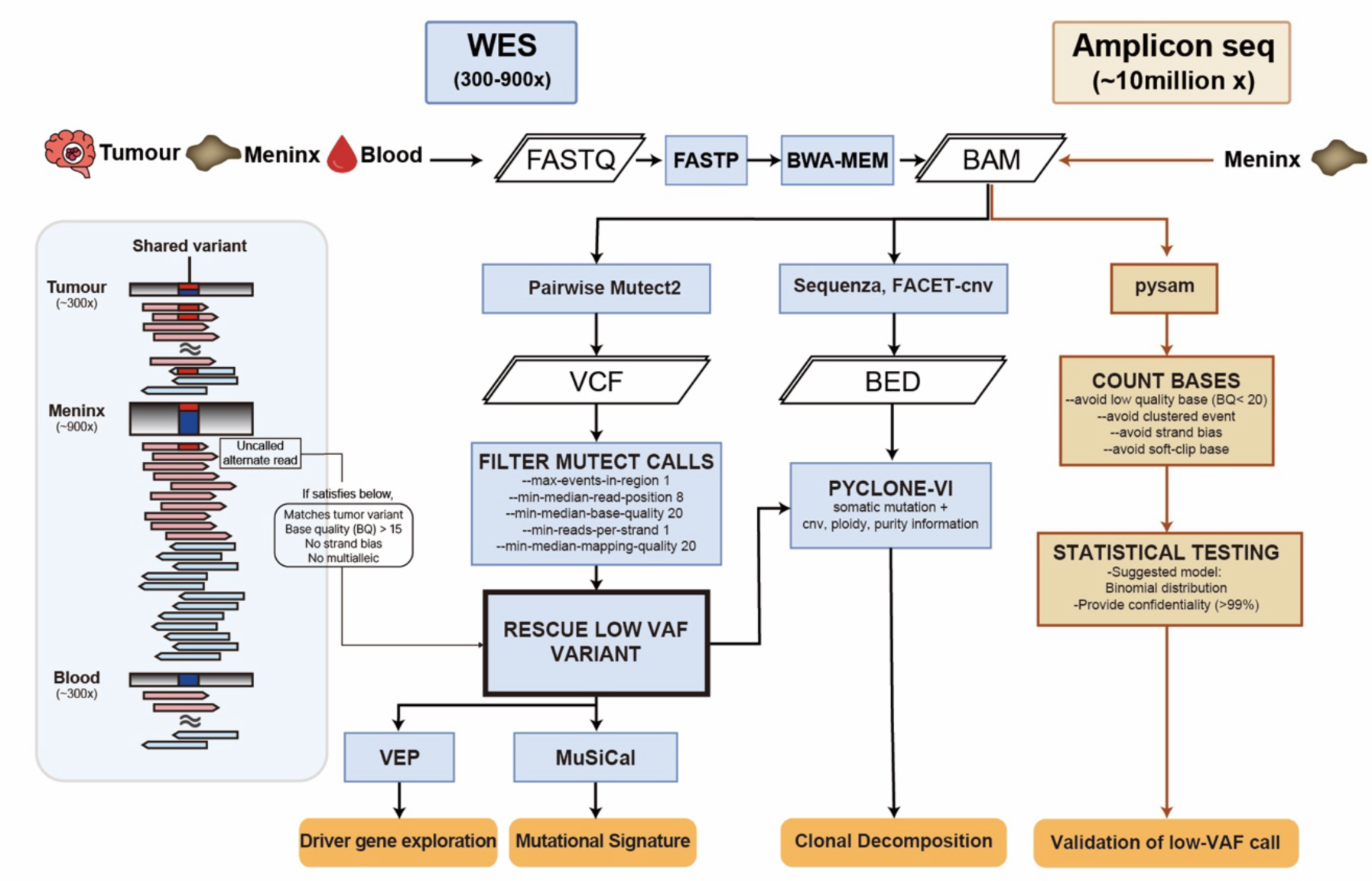
Bioinformatic pipeline for detecting shared mosaic variants. Matched samples from the tumour, meninges (dura/falx), and blood were used for WES to identify shared mosaic variants in the tumour and meninges. To identify low-allele-frequency variants in the meninges that are readily overlooked by conventional variant callers, we screened for tumour-matched candidate variants in meningeal samples using a read-level filtering strategy, retaining only high-quality evidence (left blue panel). Driver mutations in the meninges were further validated through ultra-high depth (∼10 million ×) amplicon sequencing with base-level counting and statistical testing. Downstream analyses included mutational signature inference and clonal decomposition using integrated SNV/CNV information. WES, whole-exome sequencing; VAF, variant allele frequency.

**Extended Data Fig. 6.**
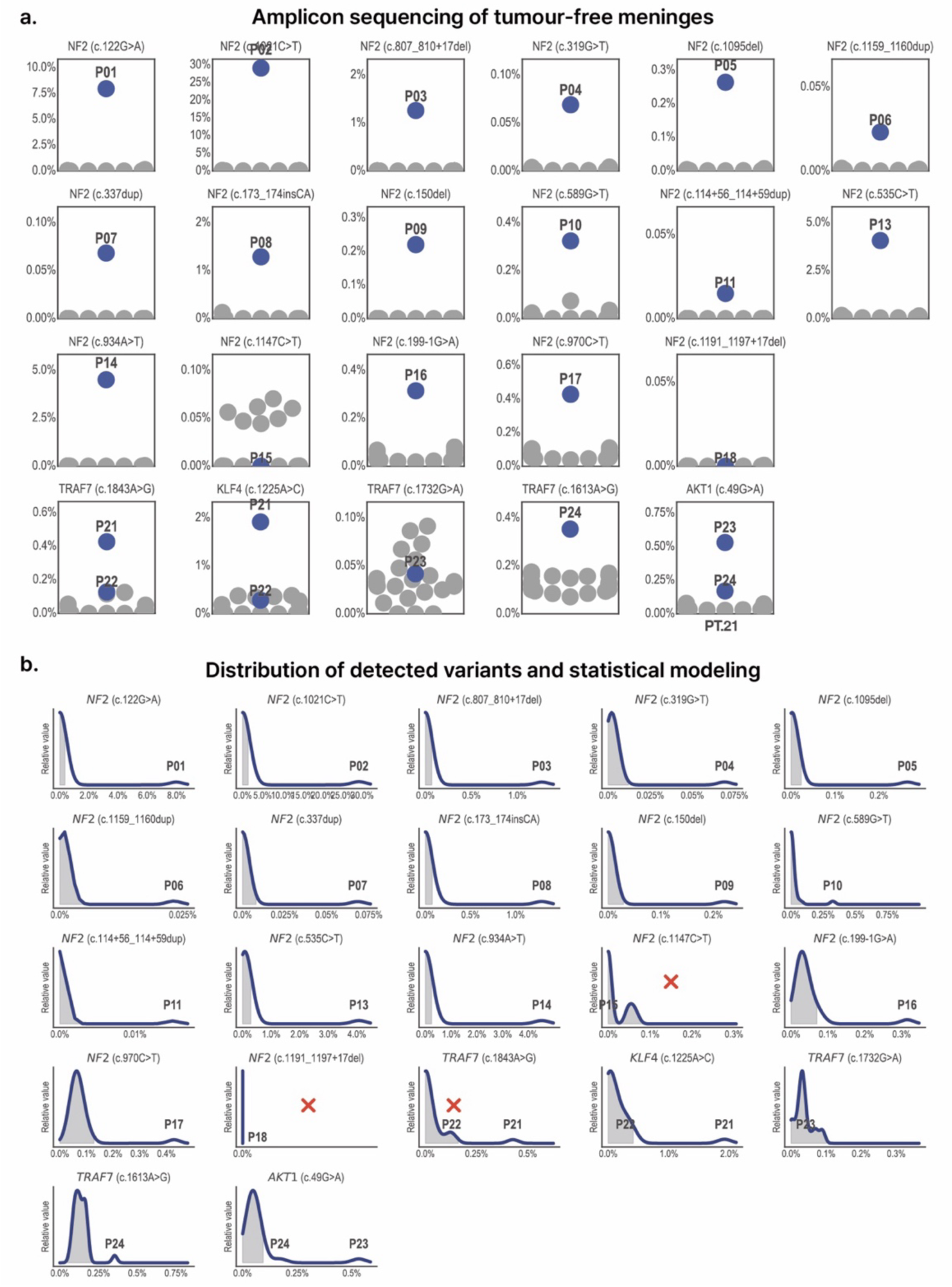
Statistical validation of variants detected by amplicon sequencing. **a,** Ultra-high-depth (∼10 million ×) amplicon sequencing was performed across all available meningeal samples at each tumour-defined driver locus. Individuals displaying driver mutations in the tumour are marked with blue circles. In some genomic positions, very low-allele frequency alternate reads were detected across all samples, necessitating careful differentiation from sequencing artefacts/background errors. **b,** VAF distribution of observed variants at each locus. The grey background represents the 99.9% confidence interval for sequencing artefacts, modelled by a binomial distribution based on the observed background error rate. Blue points indicate the patient-of-origin tumour variant, and red crosses indicate loci that did not pass the statistical threshold for calling a shared low-VAF variant in meninges (**Methods**). VAF, variant allele frequency

**Extended Data Fig. 7.**
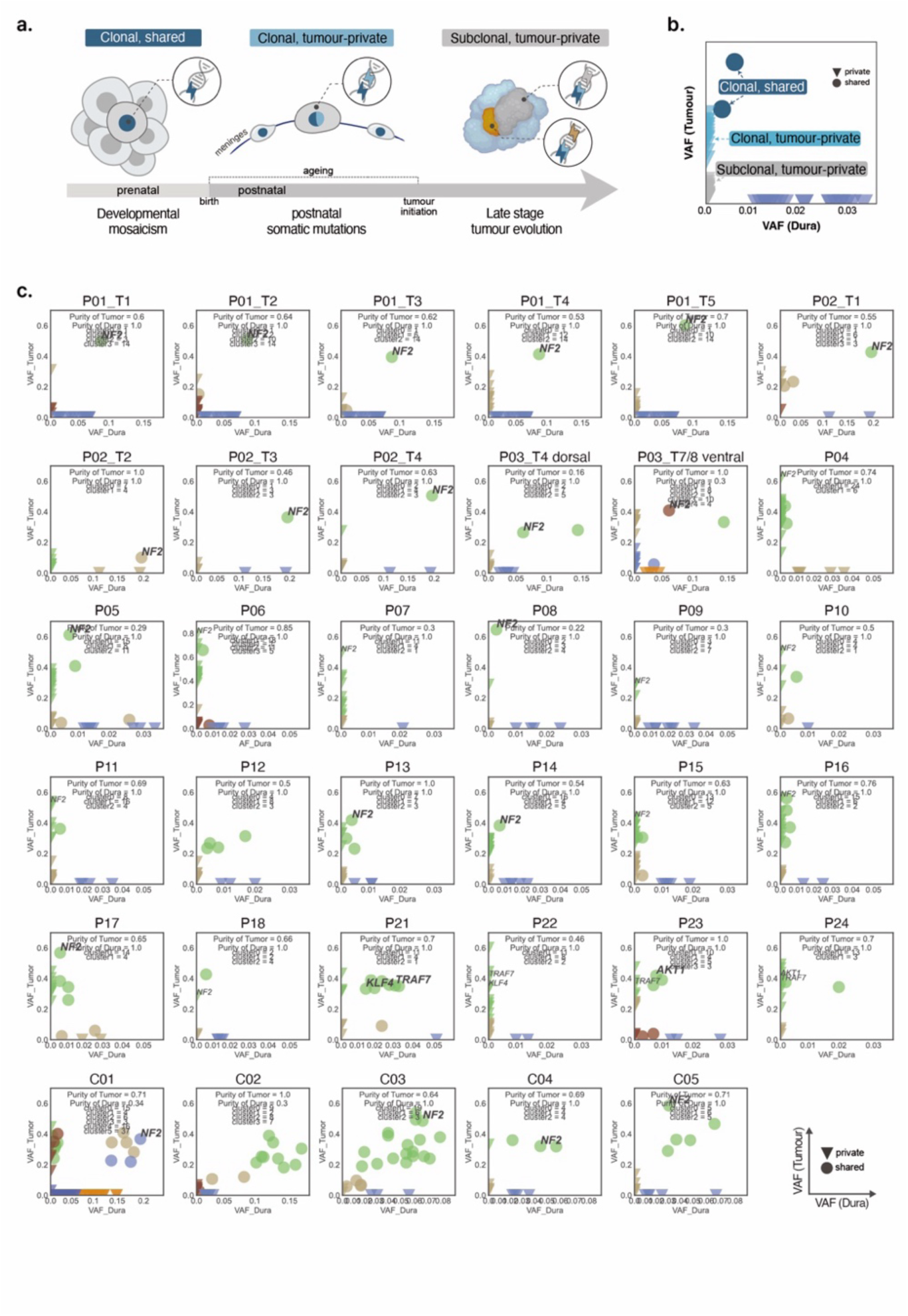
Clonal decomposition of mosaic variants. **a,** Schematic illustration of mutation acquisition timing inferred from mutational profiles in tumours and their matched meninges (dura/falx). The distribution of clonal shared, clonal tumour-private, and subclonal tumour-private mutations enables reconstruction of a temporal sequence along development, linking early developmental mosaicism, postnatal somatic mutations, and late-stage tumour evolution. **b,** Representative examples of two-sample clonality decomposition utilising matched tumour and normal meningeal pairs. Clusters are categorised as clonal shared, clonal tumour-private, or subclonal tumour-private based on their presence and clonal fractions across the matched samples. **c,** Comprehensive two-sample decomposition results for all tumour–meningeal pairs within the cohort. Driver mutations identified in the tumours are annotated, with variants also detected in the matched normal meninges highlighted in bold. VAF, variant allele frequency

**Extended Data Fig. 8.**
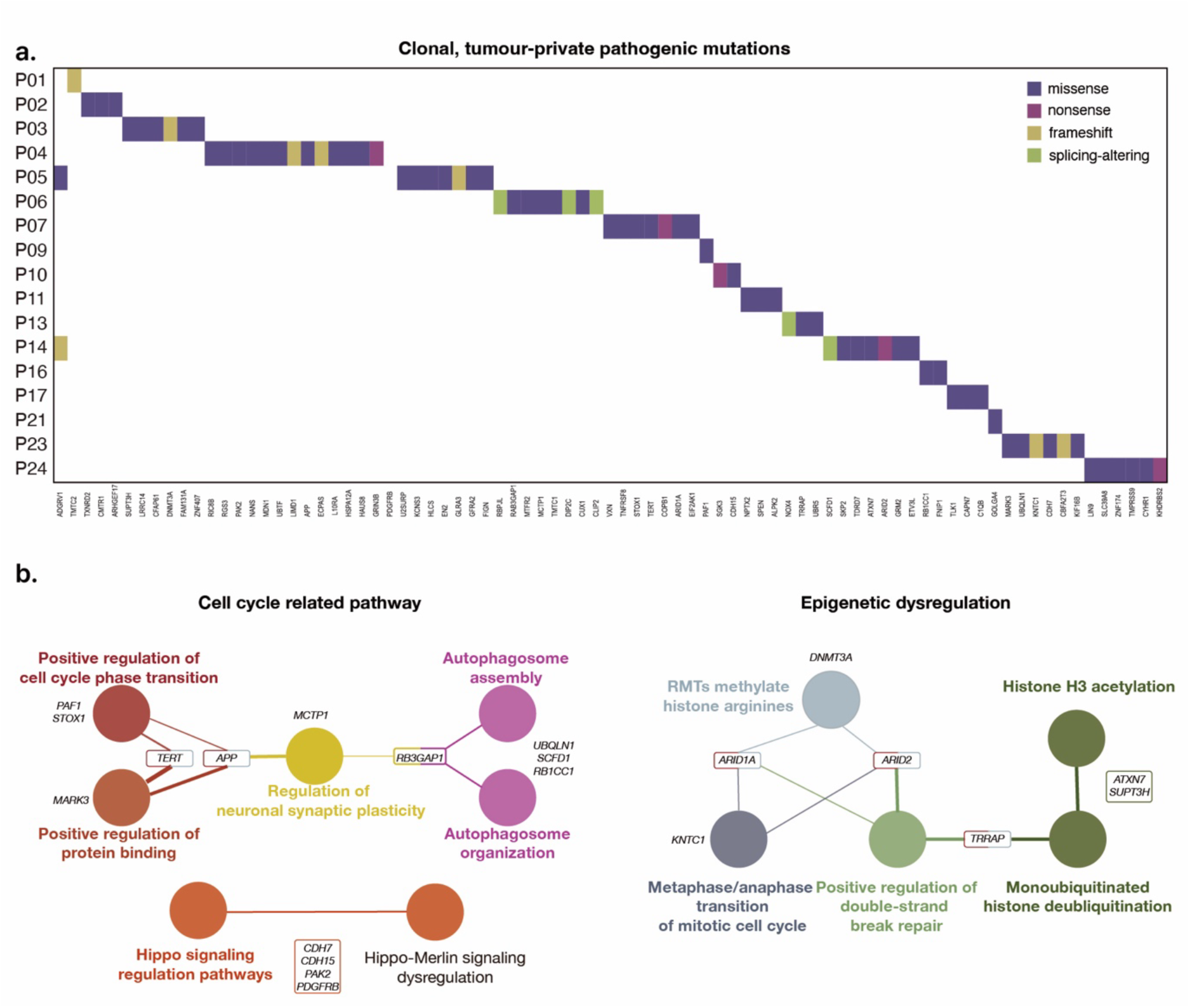
Potential tumour-driving mutations. **a,** Heatmap of pathogenic clonal mutations identified in tumour tissue, excluding *NF2*, *TRAF7*, *KLF4*, and *AKT1*, and not detected in the matched dura. **b,** ClueGO diagram showing enriched biological terms (GO, KEGG, and WikiPathways) associated with these clonal mutations, potentially contributing to tumourigenesis following early two-hit *NF2* pathway disruption and subsequent tumour evolution.

**Extended Data Fig. 9.**
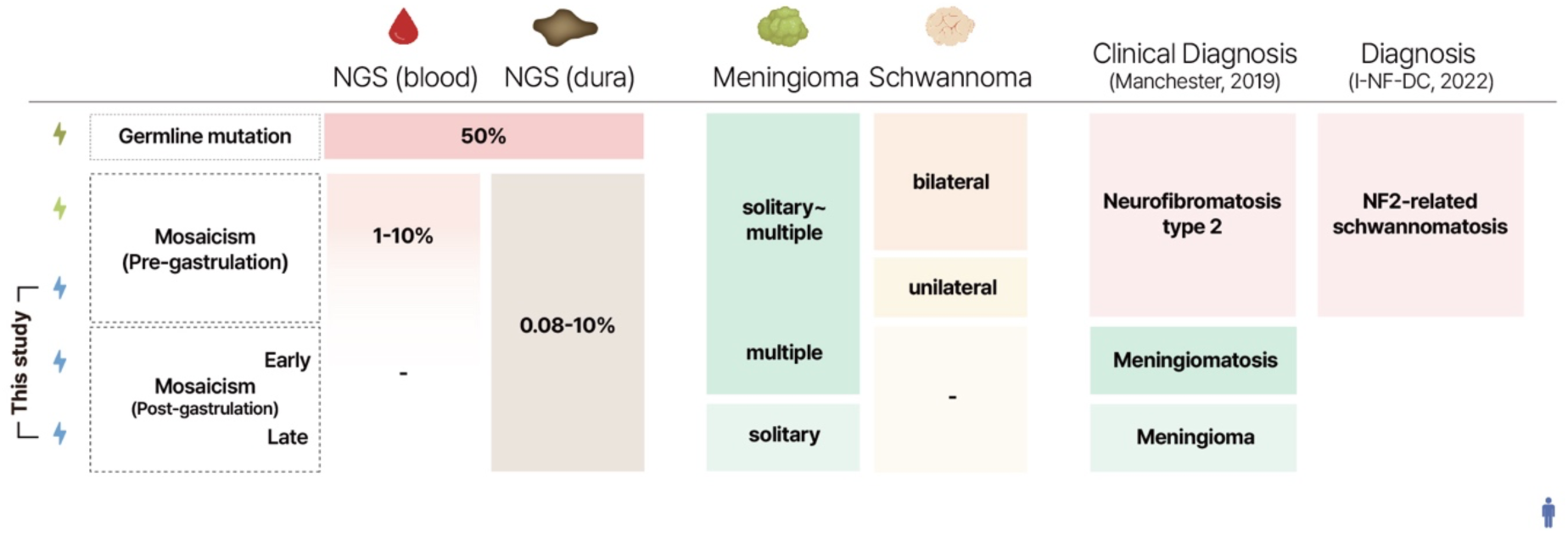
Low allele-frequency developmental mosaicism unveils the broader spectrum of *NF2*-related diseases. A schematic diagram illustrating the relationships among genetic mechanisms, phenotypes, and clinical diagnoses. Current diagnoses rely on clinical phenotypes, such as tumour type and multiplicity, as defined in the Revised Version of Manchester criteria (2019)^12^ or I-NF-DC (2022)^13^, partially assisted by genetic testing of peripheral blood. This schematic highlights how developmental mosaicism, including later lineage-restricted mosaicism, may underlie meningioma and other *NF2*-related diseases. By enabling detection of low-allele-frequency mosaic variants, this framework suggests that the spectrum from *NF2*-associated syndromic disorders to solitary meningioma may be shaped by the developmental timing and spatial distribution of driver mutations. I-NF-DC; International Consensus Group on Neurofibromatosis Diagnostic Criteria.

**Extended Data Fig. 10.**
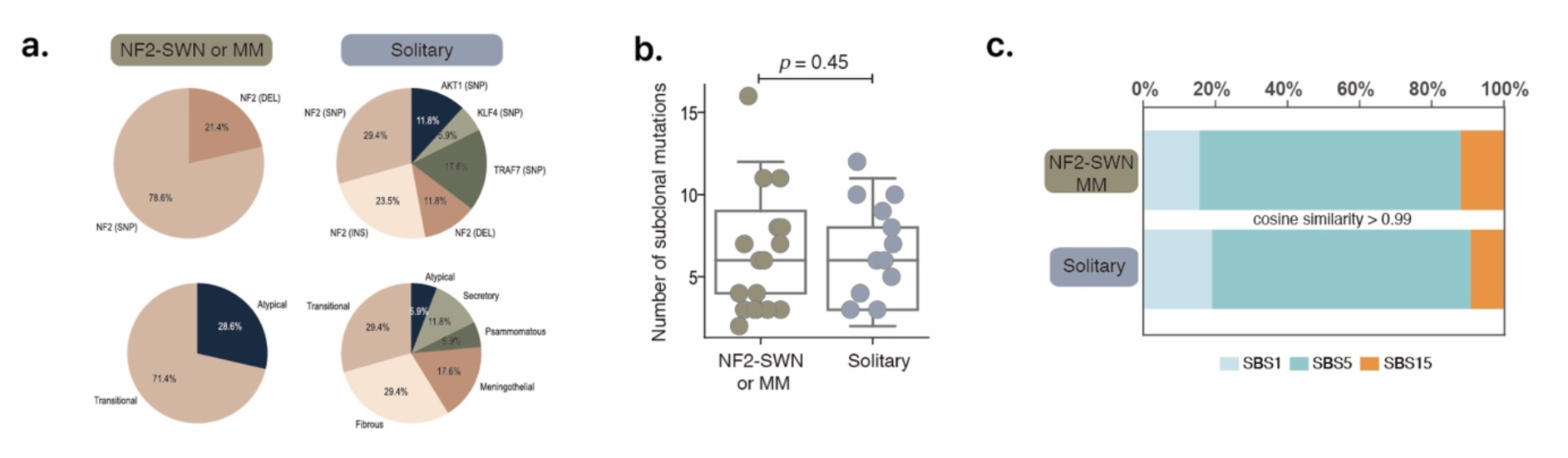
Comparable genetic architectures of solitary and multiple meningiomas. **a,** Tumour subclonal mutational burden, quantified as the number of subclonal mutations per tumour, shows no significant difference between solitary and multiple meningiomas (mean ± s.d.; two-sided Wilcoxon rank-sum test). **b,** The distribution of driver mutations and histological diagnoses does not differ substantially between the two groups. **c,** Mutational signature composition is comparable, with no evidence of differential aetiological processes as assessed by SBS contributions.

## Acknowledgement

We thank MID (Medical Illustration & Design), as a member of the Medical Research Support Services of Yonsei University College of Medicine, providing excellent support with the medical illustration. We also thank InDNA, JSLink Inc., TheragenBio, Corp., and Macrogen, Inc. for their contributions to library preparation, sequencing, and sequencing data generation. Figure 4d and 5a were created with BioRender.com. S. Kim and S-G. Kang were supported by a grant of the Korea Health Technology R&D Project through the Korea Health Industry Development Institute (KHIDI), funded by the Ministry of Health & Welfare, Republic of Korea (grant number: RS-2024-00438443), Pioneer Research Center Program, Bio&Medical Technology Development Program through the National Research Foundation of Korea funded by the Ministry of Science, ICT & Future Planning (grant number: NRF2022M3C1A309202211, RS-2024-00437820, RS-2024-00408191 and RS-2025-00523374). Y. Chung was supported by a grant of the MD-PhD/Medical Scientist Training Program and Global Medical Scientist Research Program (grant number: RS-2025-02215670) through the Korea Health Industry Development Institute (KHIDI) funded by the Ministry of Health & Welfare, Republic of Korea.

